# Combining DNA and protein alignments to improve genome annotation with LiftOn

**DOI:** 10.1101/2024.05.16.593026

**Authors:** Kuan-Hao Chao, Jakob M. Heinz, Celine Hoh, Alan Mao, Alaina Shumate, Mihaela Pertea, Steven L Salzberg

## Abstract

As the number and variety of assembled genomes continues to grow, the number of annotated genomes is falling behind, particularly for eukaryotes. DNA-based mapping tools help to address this challenge, but they are only able to transfer annotation between closely-related species. Here we introduce LiftOn, a homology-based software tool that integrates DNA and protein alignments to enhance the accuracy of genome-scale annotation and to allow mapping between relatively distant species. LiftOn’s protein-centric algorithm considers both types of alignments, chooses optimal open reading frames, resolves overlapping gene loci, and finds additional gene copies where they exist. LiftOn can reliably transfer annotation between genomes representing members of the same species, as we demonstrate on human, mouse, honey bee, rice, and *Arabidopsis thaliana*. It can further map annotation effectively across species pairs as far apart as mouse and rat or *Drosophila melanogaster* and *D. erecta*.

## 1. Introduction

Recent advancements in sequencing technologies have resulted in an exponential increase in available genome assemblies. Long-read sequencing technology, which has become faster, more cost-effective, and more accurate over the past decade ^1, 2, 3, 4, 5^, has notably enhanced the accuracy and contiguity of genome assemblies. A prominent example is the telomere-to-telomere (T2T) assembly of the human genome CHM13 ^6^, which has subsequently been used to guide the assembly of multiple additional human genomes ^7, 8, 9^. As of December 2023, the NCBI database contained 30,530 eukaryotic genomes and 567,228 prokaryotic genomes ^10, 11, 12^.

Genome annotation – identifying genes and other biological features – is essential to understanding the biology of these genomes ^13^. Annotating eukaryotic genomes presents particular challenges ^14, 15^ due to their multi-exon gene structures, extensive intergenic regions, and sparse gene density. This complexity necessitates detailed analysis and manual curation, making the process slower and more difficult to automate than genome assembly ^14, 16, 17, 18^. One widely used approach to annotation is *ab initio* gene prediction, but even the best *ab initio* systems miss many genes and struggle to get exon-intron structures precisely correct ^19, 20, 21, 22, 23, 24, 25^. RNA sequencing can more accurately identify genes by directly capturing gene transcripts, although it may miss genes if they only expressed at low levels or in certain hard-to-capture tissues ^26, 27^. Annotation "lift-over" strategies, which transfer annotations from a well-annotated genome to a newly sequenced one based on homology and synteny, provide a more efficient and cost-effective annotation method, particularly when annotations from the same or closely related species are available.

Currently, the best approach for transferring annotation between assemblies is DNA-based, exemplified by the Liftoff ^28^, and CAT ^29^ programs, which were used to create the initial annotation of the human T2T-CHM13 genome ^6^ based on the annotation of the human reference genome, GRCh38. However, in cases where the DNA sequence of a newly assembled genome deviates substantially from the reference genome, a DNA-based alignment process sometimes produces transcripts with invalid open reading frames or with erroneous splice sites. DNA-based mapping becomes even more challenging when the new genome is less closely related to the available reference.

By comparison with DNA, protein sequences of orthologous genes are conserved at much greater evolutionary distances. This observation motivated us to develop a new method that integrates protein sequence alignment into the lift-over process. A key step in this approach is to align protein sequences from the reference genome to the target, considering all six reading frames and allowing for spliced alignments to span introns. However, protein-based alignment alone cannot map all annotation between two genomes. First and most obviously, it cannot capture untranslated regions (UTRs) on either end of a transcript. Second, when small exons are separated by much longer introns, as is common in the human genome, protein-based aligners may miss some small exons entirely (as illustrated in Figures S1A-B). Third, because intronic alignments are not considered, protein-to-DNA alignment is susceptible to aligning proteins to pseudogenes. Fourth, this approach sometimes combines coding sequences (CDSs) from distinct genes when multiple members of a gene family are arranged in tandem along a genome (Figure S1C). Finally, another obvious limitation is that a protein alignment strategy cannot transfer annotations of non-coding genes or other features.

LiftOn is a homology-based genome annotation tool that uses both DNA and protein sequence alignment, and that builds on Liftoff, the current leading homology-based annotation lift-over tool. LiftOn includes several key functions to create annotation: (1) it uses a protein-maximization algorithm that combines both DNA and protein sequence alignment to generate protein-coding gene annotations that maximize similarity to the reference proteins; (2) it checks alternative open reading frames (ORFs) for truncated proteins to identify the ORF that yields the longest match to the reference protein; (3) similarly to Liftoff, it finds extra copies of protein-coding gene copies in the target genome; (4) it reports the various types of mutations for proteins that fail to match the reference perfectly, similarly to LiftoffTools ^30^; and (5) it resolves issues such as overlapping gene loci and multi-mapping for genes within a large gene family. By combining the advantages of DNA and protein sequence alignment, LiftOn generates better protein-coding gene annotation than either alignment method can achieve on its own.

In our experiments below, we demonstrate how LiftOn can yield substantial improvements in mapping annotations from one human genome, GRCh38, to another, T2T-CHM13, using three different human annotation sets: MANE ^31^, CHESS ^32^, and RefSeq ^33^. Additionally, we show that LiftOn is effective for mapping annotation between members of non-human species, including *Mus musculus* (mouse), *Apis mellifera* (honey bee), *Arabidopsis thaliana* (thale cress), and *Oryza sativa* (rice). To demonstrate LiftOn’s effectiveness at mapping annotation between distinct but closely related species, we mapped human genes onto *Pan troglodytes* (chimpanzee). Finally, we illustrate that LiftOn works on more distantly related species by mapping annotation from *Drosophila melanogaster* to *Drosophila erecta* and from *Mus musculus* to *Rattus norvegicus*.

## 2. Results

### 2.1 A two-step protein-maximization algorithm to improve protein-coding gene annotations

LiftOn implements a two-step protein-maximization (PM) algorithm (illustrated in Figure 1 and discussed in more detail in Methods) to find the best annotations at protein-coding gene loci.

**Figure 1.**
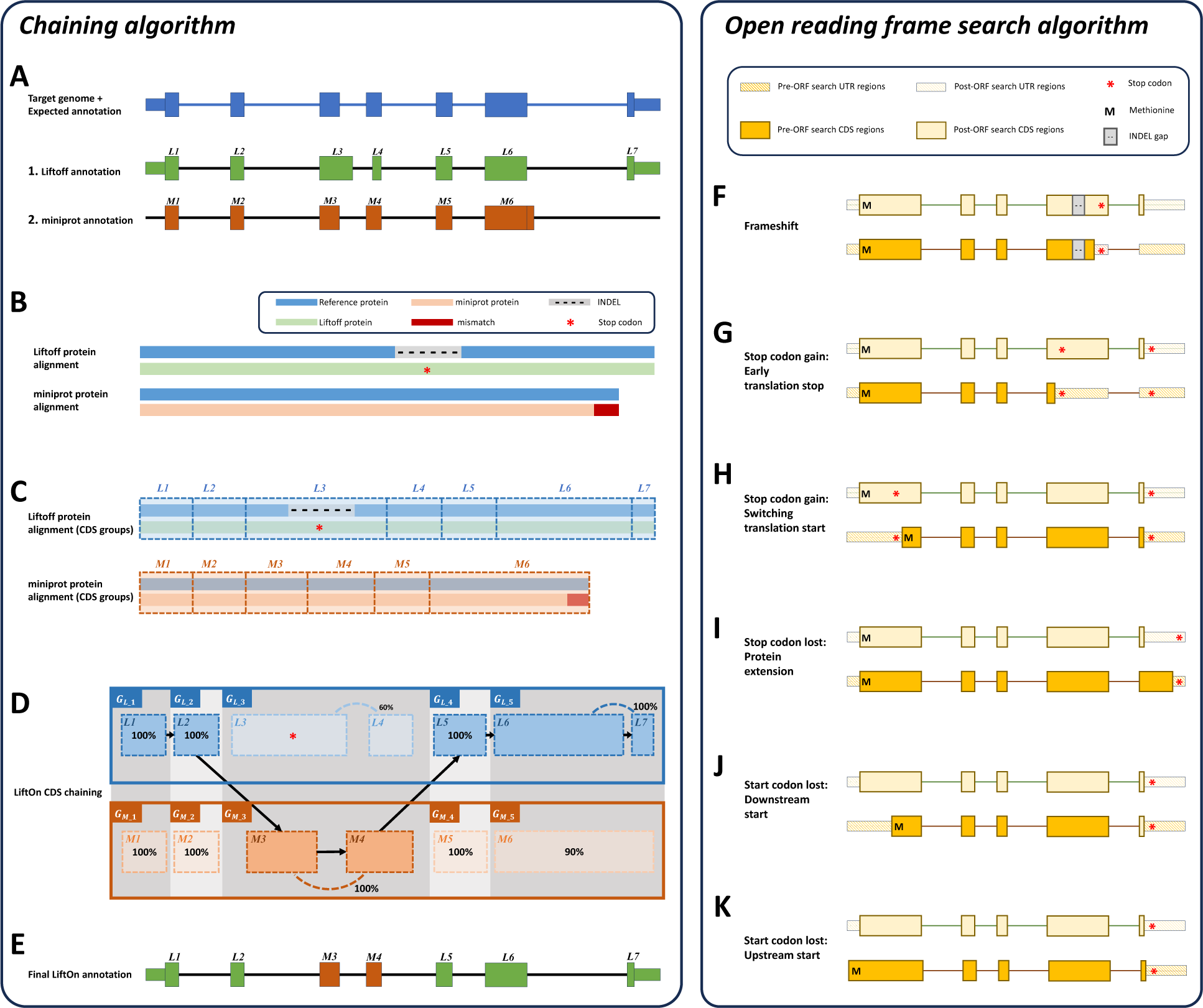
The protein-maximization (PM) algorithm consists of two modules: **(A-E)** the chaining algorithm, and **(F-K)** the open reading frame search algorithm. **(A)** Matched protein-coding transcripts mapped by Liftoff (green) and miniprot (orange) at the same location in a target genome. The transcript in blue represents the correct transcript annotation on the target genome. Liftoff’s mapping has an erroneous splice junction between L3 and L4, while miniprot’s mapping has a missing splice junction in M6. **(B)** Pairwise alignment results of the proteins mapped by Liftoff and miniprot to the reference protein. The figure shows a premature stop codon in the Liftoff protein, while the miniprot alignment has a mismatched protein sequence at the end. **(C)** Pairwise alignment mappings with added exon/CDS boundaries. **(D)** CDSs are grouped based on the cumulative lengths of the amino acids in the reference protein as described in the main text. In this example, CDSs are organized into groups: *G*_*L*_1__ = {*L*1}, *G*_*M*_1__ = {*M*1}, *G*_*L*_2__ = {*L*2}, *G*_*M*_2__ = {*M*2}, *G*_*L*_3__ = {*L*3, *L*4}, *G*_*M*_3__ = {*M*3, *M*4}, *G*_*L*_4__ = {*L*5}, *G*_*M*_4__ = {*M*5}, *G*_*L*_5__ = {*L*6, *L*7} and *G*_*M*_5__ = {*M*6}. The chaining algorithm iterates through each group, comparing the corresponding partial protein sequences to the reference protein and chaining those with higher protein sequence identity. **(E)** In this example, *G*_*L*_1__, *G*_*L*_2__, *G*_*M*_3__, *G*_*L*_4__ and *G*_*L*_5__ are chained, forming the new protein-coding transcript CDS list. This list includes *L*1, *L*2, *M*3, *M*4, *L*5, *L*6 and *L*7 in the LiftOn annotation. **(F-K)** Schematic diagrams illustrating how the ORF search algorithm handles various types of sequence mutations. This process leads to changes in the gene annotation of both translated and untranslated regions. **(F)** Frameshift mutation: a variation caused by the insertion or deletion of a sequence of nucleotides whose length is not divisible by three. In this example, the indel introduces a premature stop codon. **(G-H)** Point mutations leading to premature stop codons. LiftOn searches for the longest ORF, considering two scenarios: **(G)** depicts the selection of the first encountered stop codon, while **(H)** illustrates the switch to the downstream start codon. **(I)** Stop codon loss: when a stop codon is deleted, LiftOn identifies a new stop codon in the 3’ UTR. **(J-K)** Start codon loss: in this scenario, LiftOn searches for a new start codon, exploring both downstream in the coding region **(J)** and upstream in the 5’ UTR **(K)**.

First, it uses a chaining algorithm, described below, to find the exon-intron boundaries of protein coding transcripts. Second, if the coding sequence (CDS) does not preserve the entire protein sequence of the reference protein, it adjusts the CDS boundaries to preserve as much of the reference protein as possible. (Note that CDS features are defined as the coding portions of exons, from start to stop, and excluding untranslated portions of exons.)

The chaining algorithm (Figure 1A-E) starts by pairing up miniprot alignments with transcripts identified by Liftoff (see Methods and Algorithm S1 for details on the pairing approach). After two transcripts are paired, the protein sequences from the Liftoff and miniprot annotations are then aligned to the full-length reference protein, as illustrated in Figure 1B. Subsequently, LiftOn maps the CDS boundaries from both the Liftoff and miniprot annotations onto the protein alignment (Figure 1C and Algorithm S2).

The CDSs within the Liftoff and miniprot annotations are grouped in the 5’ to 3’ direction. The CDSs groups are represented as *G*_*L*_*i*_ and *G*_*M*_*i*_ respectively for LiftOff and miniprot, where *i* denotes the *i^th^* group in that annotation (Figure 1D).

The grouping process begins with the first CDS in each annotation and continues until reaching the endpoints of the downstream CDS in Liftoff and miniprot, where the number of aligned amino acids from the reference protein is equal. This forms the first CDS group in Liftoff, denoted as *G*_*L*_1_, and the first CDS group in miniprot, denoted as *G*_*M*_1_. Subsequent groups start from the previous endpoint in both Liftoff and miniprot, extending until the number of aligned amino acids from the reference protein matches for both annotations again. These subsequent groups are represented as *G*_*L*_2_ and *G*_*M*_2_, respectively. The grouping process concludes upon reaching the last CDS in both annotations (see Algorithm S3 for more details).

Within each group, *G*_*L*_*i*_ or *G*_*M*_*i*_, we calculate the partial protein sequence identity and select the group with higher protein sequence identity score (Figure 1D and Algorithm S4). In case of a tie, LiftOn prioritizes the Liftoff annotation, *G*_*L*_*i*_, so that it will include UTRs in its output. The selected group of CDSs is represented by *G*_*SEL*_*i*_. All CDSs in *G*_*SEL*_*i*_ are then concatenated to form the final LiftOn transcript, as shown in Figure 1E. This transcript is an ordered sequence of CDSs sourced from either Liftoff or miniprot, with the goal of maximizing protein similarity to the reference protein. This approach is particularly effective in addressing issues such as in-frame indels or mis-splicing that may arise from misalignments as illustrated in Figure 1A.

After applying the chaining algorithm, LiftOn attempts to make further improvements in the CDS regions, as illustrated in Figure 1F-K. It searches the translations of protein-coding transcripts and adjusts CDS boundaries to avoid early stop codons (Figure 1F,G), choose better translation start sites (Figure 1H,J,K), or extend proteins with stop codon loss (Figure 1I). Figure S2 provides additional IGV ^34, 35^ screenshots that illustrate LiftOn’s results for each scenario.

After making these adjustments, LiftOn evaluates the differences between the reference and target transcripts and, similarly to LiftoffTools ^30^, produces a mutation report. Transcripts are deemed "identical" when their target and reference gene DNA sequences are entirely the same. For non-identical sequences, LiftOn categorizes their differences using these categories: synonymous, non-synonymous, in-frame insertion, in-frame deletion, frameshift, stop codon gain, stop codon lost, and start codon loss.

### 2.2 LiftOn improves human annotation lift-over

For our first experiment, we mapped annotations from GRCh38 onto T2T-CHM13 using RefSeq ^33^ (release 220) as the reference annotation (Figure S3). In total, there were 37,986 protein-coding and non-coding genes, with 160,561 transcripts in the reference annotation, of which 130,528 were protein-coding (see Table S1 for all mapped genes and Methods 4.7 for our gene counting approach). We focused our analysis on the protein-coding transcripts because those are the ones where LiftOn can produce improvements over DNA-based methods. At the gene level, LiftOn successfully lifted over 37,828 genes, while 158 genes remained unmapped, of which 103 were protein-coding. The overall gene mapping rate was 99.6%. Out of the successfully mapped genes, 37,453 were mapped as single copies and 375 genes were mapped with extra copies. In total, 38,886 gene loci were mapped onto T2T-CHM13 (Table 1).

**Table 1.** Statistics for LiftOn at both the gene and transcript levels, as a result of mapping RefSeq release 220 annotation from the GRCh38 human genome to T2T-CHM13.

|  |  | Total feature count | Protein-coding feature count |  |  | Non-coding feature count |  |  |
| --- | --- | --- | --- | --- | --- | --- | --- | --- |
| Gene | Reference (GRCh38) | 37,986 | 19,927 |  |  | 18,059 |  |  |
|  | Target (CHM13) | 38,916 | Single copy | >1 copy | Extra copy | Single copy | >1 copy | Extra copy |
|  |  |  | 19,738 | 86 | 320 | 17,715 | 289 | 768 |
|  |  |  | 20,144 |  |  | 18,772 |  |  |
| Transcript | Reference (GRCh38) | 160,561 | 130,528 |  |  | 30,033 |  |  |
|  | Target (CHM13) | 16,1701 | Single copy | >1 copy | Extra copy | Single copy | >1 copy | Extra copy |
|  |  |  | 129,967 | 239 | 573 | 29,488 | 410 | 1,024 |
|  |  |  | 130,779 |  |  | 30,922 |  |  |

In assessing overall protein-coding gene annotations, we computed gap-compressed protein sequence identity scores for LiftOn, Liftoff, and miniprot. Figure 2A compares the mapping of protein-coding genes using Liftoff, which is based on DNA alignment, to miniprot, which uses protein-to-DNA alignment. Dots in the lower right part of the panel indicate transcripts where Liftoff was superior to miniprot, as measured by protein sequence identity, while the upper left part of the plot shows transcripts where miniprot was superior. This comparison shows that neither approach dominates the other. In the LiftOn versus Liftoff comparison (Figure 2B), 866 transcripts exhibit higher protein sequence identity, with 113 of those achieving 100% identity. Similarly, the LiftOn versus miniprot comparison (Figure 1C) shows that LiftOn finds better matches for 30,266 protein-coding transcripts, improving 22,746 to 100% identity. The protein sequence identity distributions are shown in Figure S4.

**Figure 2.**
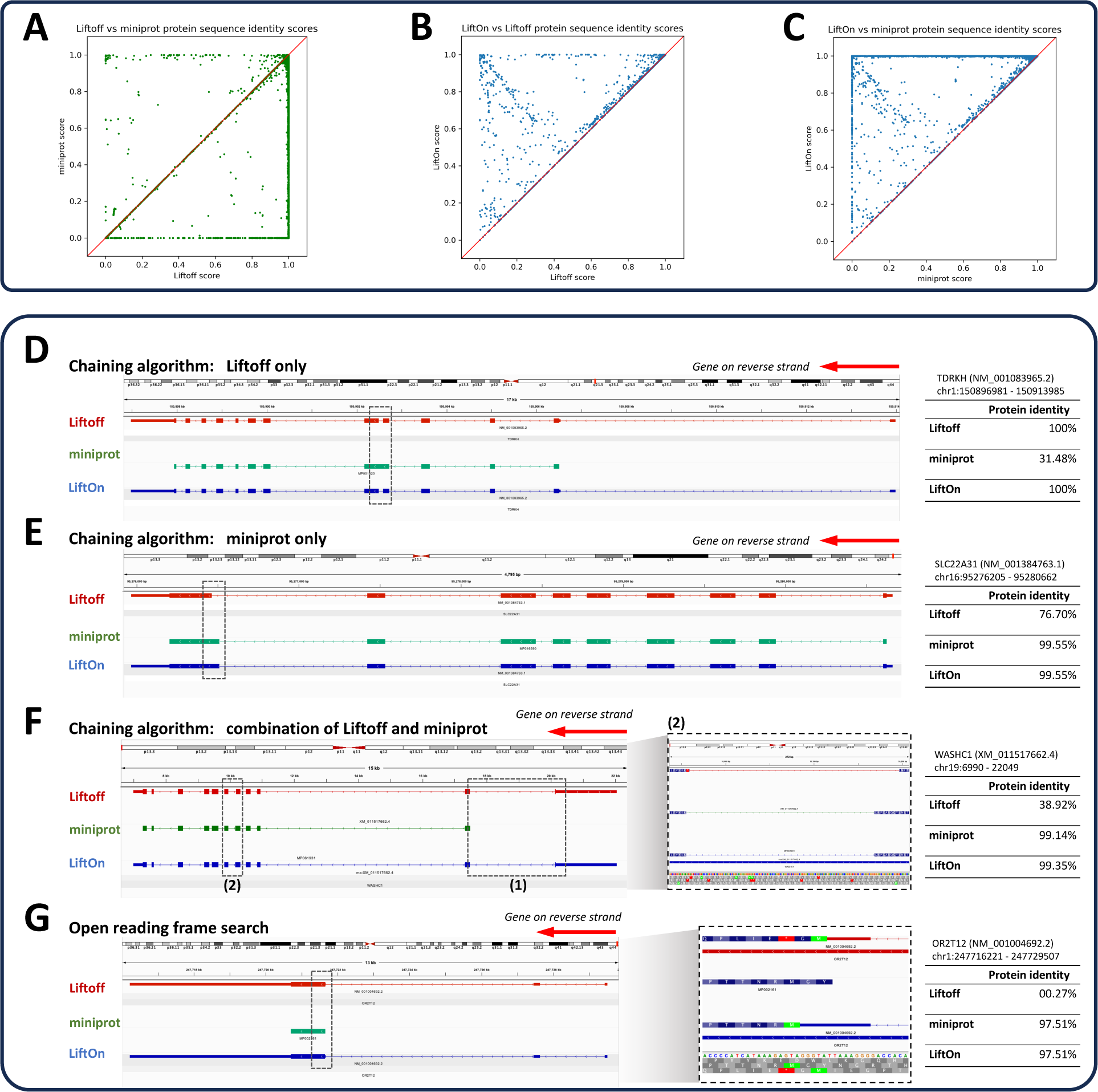
Comparative analysis of various tools for mapping RefSeq protein annotations from GRCh38 to T2T-CHM13 v2.0. **(A-C)** Scatter plots of protein sequence identity. **(A)** Comparison between miniprot (y-axis) and Liftoff (x-axis), **(B)** comparison between LiftOn (y-axis) and Liftoff (x-axis), and **(C)** comparison between LiftOn (y-axis) and miniprot (x-axis). **(D-G)** Examples of improved annotation due to LiftOn’s two-step PM algorithm. **(D)** LiftOn employs Liftoff’s annotation to correct a splice junction missed by miniprot in transcript NM_001083965.2 of the TDRKH gene. **(E)** For transcript NM_001384763.1 of the SLC22A31 gene, LiftOn uses miniprot’s annotation to resolve an incorrect acceptor site from Liftoff’s annotation. **(F)** For transcript XM_011517662.4 of the WASHC1 gene, LiftOn combines both annotations to rectify an omitted CDS by miniprot and a misidentified splice junction by Liftoff between the fifth and sixth exons. **(G)** LiftOn’s ORF search algorithm selects an alternative downstream start codon for a frameshift mutation in transcript NM_001004692.2 of the OR2T12 gene, thereby conserving the majority of the protein sequence.

The LiftOn PM algorithm can improve protein-coding transcript annotations in four scenarios illustrated in Figure 2. First, as observed with the TDRKH gene (Figures 2D and S5A), miniprot missed a splice junction, leading to the retention of an intron and a premature stop codon. Here LiftOn adopted Liftoff’s CDS, which yielded the correct protein. Second, as shown in Figures 2E and S5B for the SLC22A31 gene, miniprot correctly identified an AG acceptor site that Liftoff missed, yielding the correct CDS, which LiftOn combined with the UTRs from Liftoff. Third, LiftOn sometimes was able to fix errors in both methods, as shown in Figures 2F and S5C for the gene WASHC1. In this example, LiftOn corrected the omission of the first coding exon by miniprot and fixed an incorrect splice junction found by Liftoff, one that erroneously introduced a premature stop codon between the fifth and sixth exons. LiftOn’s chaining algorithm effectively consolidates these segments, improving the protein sequence identity to 99.4%.

Fourth, as shown in Figure 2G for the OR2T12 gene, a mutation introduced a premature stop codon (Figure 2G, inset), and the ORF search algorithm in LiftOn adjusted by scanning for ORFs that best match the reference protein sequence. Here it found an alternative start codon just two codons downstream that maintained a near full-length match to the reference protein.

LiftOn identifies and reports additional copies of lifted-over features, similarly to Liftoff. In the mapping from GRCh38 to CHM13, LiftOn identified 86 protein-coding genes with at least one extra copy, for a total of 320 new protein-coding gene loci. Of these, 289 extra gene copies were detected by Liftoff and 31 by miniprot, detailed in Tables S2 and S3. Figure 3 shows the relative positions of extra gene copies between the two genomes, illustrating that most were located on the same chromosomes.

**Figure 3.**
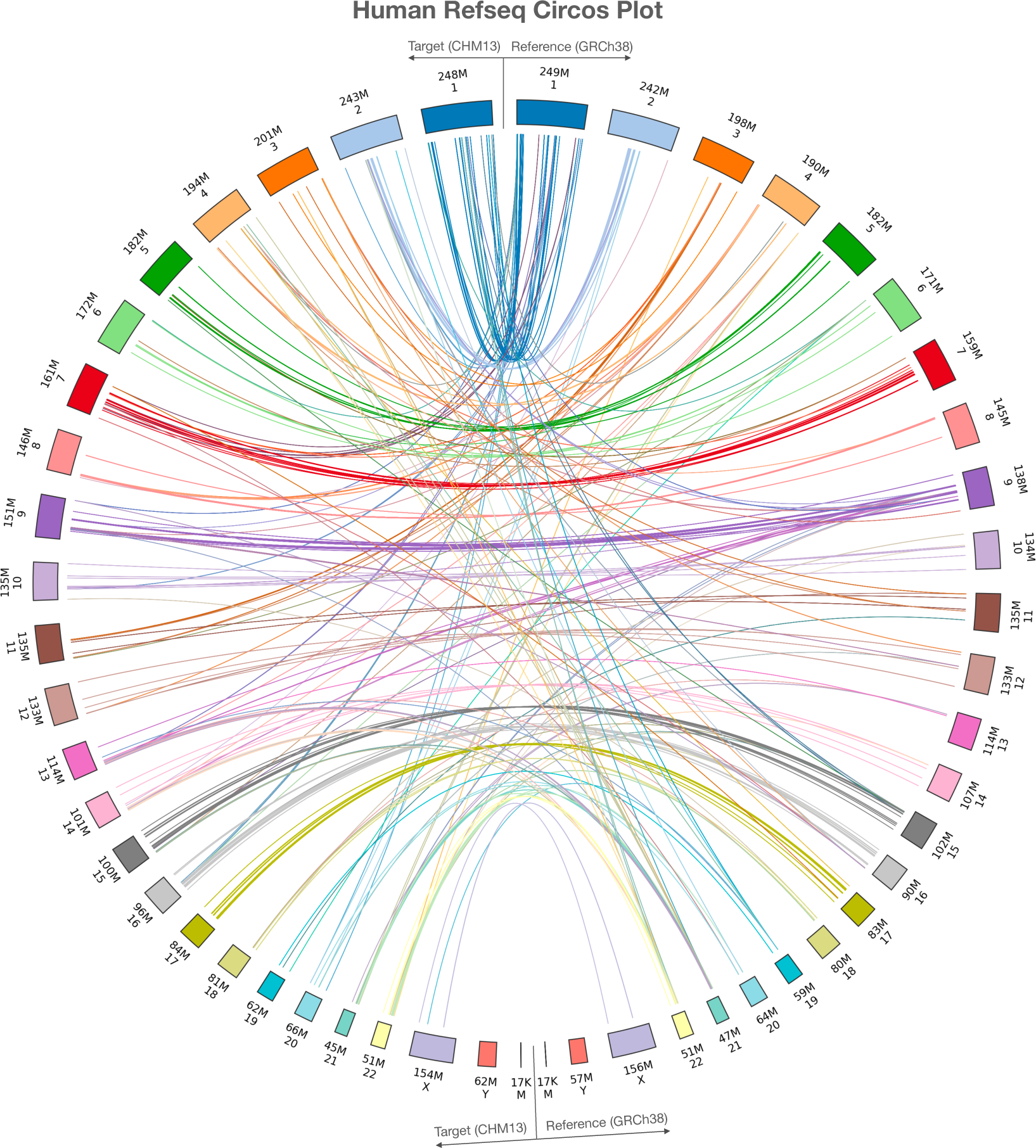
Plot illustrating the locations of extra gene copies found on T2T-CHM13 (left side) compared to GRCh38 (right side). The Circos plot were generated using pyCircos (https://github.com/ponnhide/pyCircos). Each line represents a gene copy mapped from the reference genome to the target genome, with colors indicating different chromosomes. The lines are color-coded by the chromosome of the original copy. The bands within the plot are sized proportionally to reflect the actual size of these chromosomes.

We further compared the LiftOn-generated annotation of CHM13 with the current release of the T2T-CHM13 annotation (available from the T2T github site, https://github.com/marbl/CHM13). We excluded the annotation of chromosome Y from this analysis due to the extremely complex repeat structure of that chromosome, which confounds most attempts to align genes consistently to it ^36^. Our comparison considered only protein-coding transcripts on chromosomes 1-22 and X. LiftOn’s annotation contained 665 protein-coding transcripts that had a higher protein sequence identity to the corresponding proteins on GRCh38, as compared to the current CHM13 annotation. Four examples are shown in Figure 4, each of which illustrates how LiftOn can improve the fidelity of the match between the source protein and the mapped-over version.

**Figure 4.**
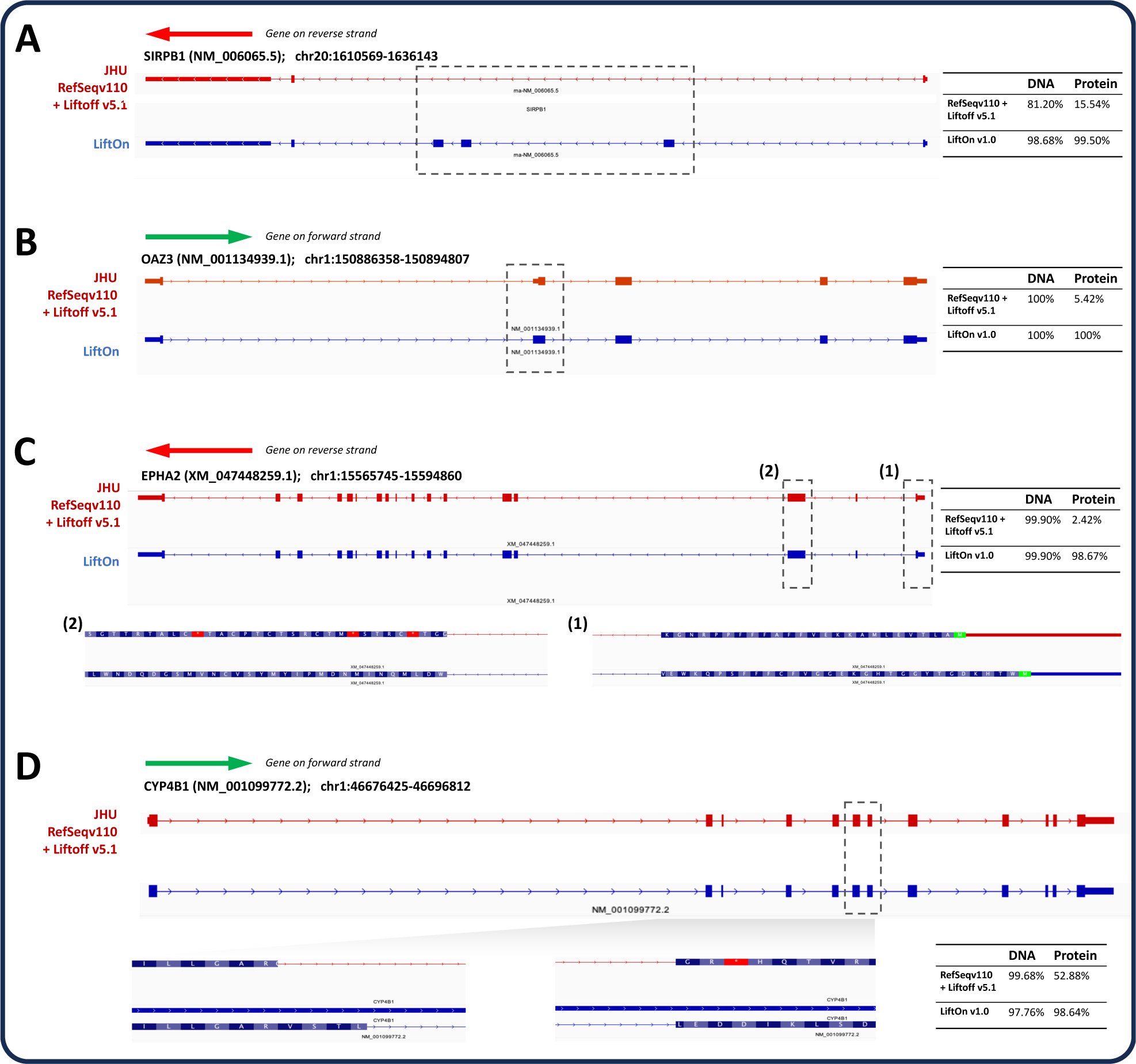
Examples where LiftOn improves the current T2T-CHM13 annotation. **(A)** In the NM_006065.5 transcript of the SIRPB1 gene, the current CHM13 annotation omits three coding exons. The LiftOn version finds those exons and increases the DNA sequence identity from 81% to 98%. **(B)** In transcript NM_001134939.1 of OAZ3, the CHM13 annotation incorporates a partial CDS in the second exon, leading to a truncated protein. LiftOn corrects this mis-annotation, increasing the protein sequence identity from 5.42% to 100%. **(C)** In transcript XM_047448259.1 of EPHA2, the published annotation chooses the wrong start codon. LiftOn finds a better start codon that improves the protein sequence identity from 2.4% to 98.7%. **(D)** In transcript NM_001099772.2 from CYP4B1, LiftOn shifts the donor site of the seventh coding exon by 11 nucleotides, fixing a frameshift and improving the protein sequence identity from 53% to 99%.

Among the 130,528 protein-coding transcripts mapped onto CHM13 by LiftOn, 13,486 transcripts were identical and 86,335 had only synonymous mutations. Of the remaining 30,707 transcripts, only 1,046 had a protein with less than 95% identity to the reference protein. Overall, LiftOn’s mapped-over annotation has greater fidelity to the source annotation (from GRCh38) than the current T2T-CHM13 annotation (Figure S6).

We observed similar improvements (as compared to both Liftoff and miniprot) when we lifted over different human annotations, including MANE ^31^ (Note S1, Table S4 and Figure S7) and CHESS3 ^32^ (Note S2, Table S5 and Figure S8). Additionally, to assess the ability of LiftOn to map annotation between genomes of non-human species, we ran experiments on two different genomes from each of four plant and animal species: *Mus musculus* ^37, 38^ (Note S3, Table S6 and Figure S9), *Apis mellifera* ^39, 40^ (honey bee, see Note S4, Table S7 and Figure S10), *Oryza sativa* ^41, 42^ (Asian rice, Note S5, Table S8 and Figure S11), and *Arabidopsis thaliana* ^43, 44^ (Note S6, Table S9 and Figure S12), and obtained comparable enhancements. These analyses demonstrate the robust performance of LiftOn in handling annotation lift-over across a broad spectrum of organisms.

### 2.3 LiftOn improves both closely and distantly related species annotation lift-over

To demonstrate that LiftOn can reliably lift-over cross-species annotations, we used it to transfer genes between both closely-related and more distantly-related species. We measured genomic distance using Dashing ^45^ and Mash ^46^.

#### 2.3.1 Mapping from Homo sapiens to Pan troglodytes

Chimpanzees are the closest evolutionary relatives of humans ^47, 48, 49, 50^, so we chose this example to demonstrate the applicability of LiftOn between closely-related species. LiftOn successfully lifted-over 37,509 genes (Table 2, single copy + >1 copy), leaving 477 genes unmapped, of which 285 were protein-coding and 192 noncoding. The overall gene mapping rate was 98.7%. Out of all mapped genes, 37,081 were mapped uniquely, while 428 genes were mapped to multiple locations, producing a total of 38,879 gene loci. LiftOn obtained higher protein sequence identity scores than Liftoff and miniprot for 4,332 and 33,509 transcripts respectively (Figure 5A, Note S7, and Figure S13).

**Figure 5.**
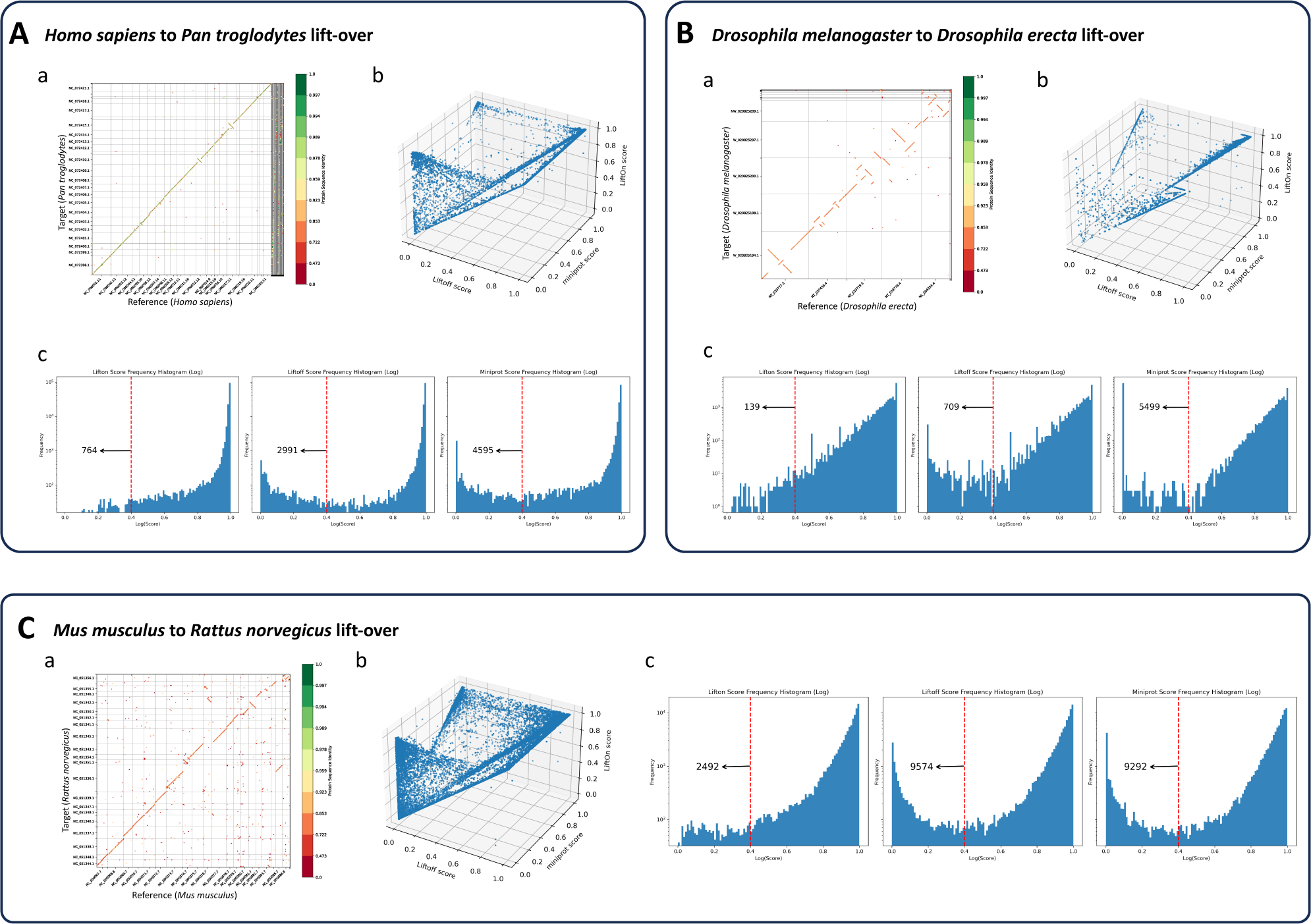
**(A)** Comparative analysis of lifting over RefSeq v220 annotations from *Homo sapiens* (GRCh38) to *Pan troglodytes* (NHGRI_mPanTro3-v1.1). **(B)** Comparative analysis of lifting over annotations from *Drosophila melanogaster* (genome assembly release 6 + ISO1MT) to *Drosophila erecta* (Prin_Dsim_3.1). **(C)** Comparative analysis of lifting over annotations from *Mus musculus* (GRCm39) to *Rattus norvegicus* (mRatBN7.2). Graphs labeled (a) show protein-gene order plots, with the x-axis representing the reference genome and the y-axis representing the target genome. The protein sequence identities are color-coded on a logarithmic scale, ranging from green (identical) to red. Graphs labeled (a) show protein-gene order plots, with the x-axis representing the reference genome and the y-axis representing the target genome. The protein sequence identities are color-coded on a logarithmic scale, ranging from green (1) to red (0), represent the degree of amino acid similarity, with 1 indicating identical sequences and 0 indicating no shared amino acids. The gene order plot script was customized from Liftofftools. Graphs labeled (b) are 3-D protein sequence identity plots comparing Liftoff on the x-axis, miniprot on the y-axis, and LiftOn on the z-axis. Each dot represents a protein-coding transcript. If a dot is above the x=y plane, LiftOn’s mapping produced a higher protein sequence identity score than the other programs. (c) Frequency plots on a logarithmic scale of protein sequence identity for LiftOn (left), Liftoff (middle), and miniprot (right).

**Table 2.**
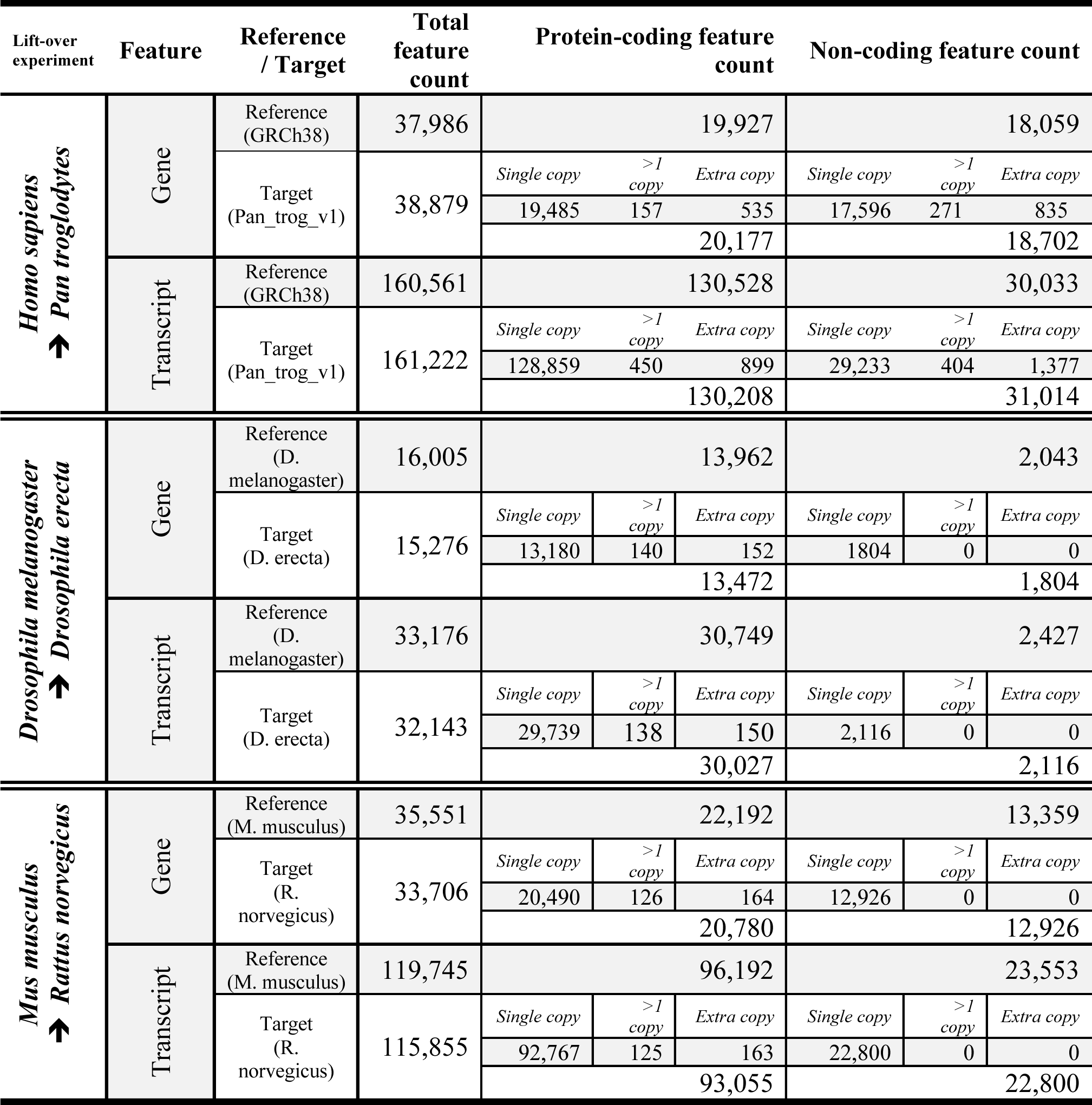
Statistics for LiftOn at both the gene and transcript levels, after mapping RefSeq release v220 annotation from the human genome to *Pan troglodytes* (mPanTro3-v1.1), from *Drosophila melanogaster* (genome assembly release 6 + ISO1MT) to *Drosophila erecta* (Prin_Dsim_3.1), and from *Mus musculus* (GRCm39) to *Rattus norvegicus* (mRatBN7.2).

#### 2.3.2 Mapping from Drosophila melanogaster to Drosophila erecta

We then mapped the annotation between two Drosophila species ^51, 52^ that are considerably more distant from each other than human and chimpanzee: *Drosophila melanogaster* ^53^ and *D. erecta* ^54, 55^, with 0.07 Dashing ^45^ similarity score and 0.08 Mash-distance ^46^. A protein-coding gene order plot (Figure 5B) shows multiple genome rearrangements, illustrating the divergence between the genomes. The LiftOn gene- and transcript-level statistics are summarized in Table 2, showing a gene mapping rate of 94.5% and a transcript mapping rate of 96.4%. LiftOn yields improved annotations for 4,583 and 7,763 protein-coding transcripts compared to Liftoff and miniprot respectively (Note S8 and Figure S14).

#### 2.3.3 Mapping from Mus musculus to Rattus norvegicus

Finally, we chose two model organisms, *Mus musculus* ^56^ and *Rattus norvegicus* ^57^ (mouse and rat), to showcase LiftOn’s ability to map annotation between even more distant species, with 0.01 Dashing ^45^ similarity score and 0.12 Mash-distance ^46^. The results are illustrated in Figure 5C and Table 2, which reveals a gene mapping rate of 94.3% and a transcript mapping rate of 96.6%. For the mouse to rat mapping, LiftOn improves the annotations for 15,420 and 30,574 protein-coding transcripts as compared to Liftoff and miniprot respectively. The frequency plots in Figure 5D reveal that LiftOn only had 2,492 protein-coding transcripts below the 40% identity threshold, a notable improvement over Liftoff (9,574) and miniprot (9,292). More details are included in Supplementary Note S9 and Figures S15-S16.

## 3. Discussion

Annotation lift-over is a powerful means to transfer annotation from one genome to another, but mapping requires accurate alignment, which increases in difficulty as genomes become more divergent. Incorporating protein sequence alignment, as we have done here, addresses some of the challenges, but factors such as pseudogenes, large gene families, structural variations, and the quality of the source annotation all affect the difficulty of the problem.

LiftOn’s accuracy will always be critically dependent on the accuracy of the source annotation. As an example, consider the mapping of two uncharacterized protein-coding genes, LOC124905331 and LOC112268317, both of which mapped onto CHM13 in multiple copies. LOC124905331 appears on an unplaced contig (NT_187388.1) in GRCh38, and on CHM13, LiftOn mapped it to chromosome 21, spanning positions 3,205,883 to 5,522,421, with 50 tandemly repeated copies. LOC112268317 was initially on another unplaced contig (NT_187499.1) in GRCh38, and LiftOn mapped it in 30 copies on CHM13, near the telomeric regions of chromosomes 14, 15, 21, and 22. Both of these genes overlap with the ribosomal DNA (rDNA) array, which occurs in many copies on the acrocentric chromosomes 13, 14, 15, 21, and 22 ^6, 9, 58^. Given this overlap with rDNA, we suspect that the source genes are not genes at all, and manual curation would be required to clean up the results of the automated lift-over process.

LiftOn is a versatile homology-based annotation mapping tool, designed as a successor to Liftoff, the leading homology-based annotation lift-over tool. LiftOn improves on Liftoff by mapping protein-coding transcripts more accurately, which it achieves through the use of a protein-to-DNA aligner, miniprot, a chaining algorithm to generate protein sequences with high similarity to reference proteins, and post-processing methods to identify and correct open reading frames that might otherwise produce truncated proteins. Our experiments demonstrate that LiftOn outperforms other methods that rely solely on DNA or proteins for mapping annotations from one genome to another. Its use of protein sequence alignment allows it to effectively map annotation between different species, even those as distantly related as mouse and rat.

## 4. Online Methods

LiftOn is implemented as a Python package that maps gene features from a reference genome to a target genome using DNA- and protein-alignment-based methods, namely Liftoff and miniprot. For inputs, LiftOn requires a reference annotation in either GFF or GTF format, as well as reference and target genomes provided as FASTA files. LiftOn uses gffutils version 0.12 (https://github.com/daler/gffutils) to create an sqlite3 database, and it uses Biopython ^59^ and Pyfaidx ^60^ to extract both DNA and protein transcript sequences from the reference genome.

Subsequently, LiftOn runs the embedded Liftoff code and runs miniprot through a python subprocess package, producing separate Liftoff and miniprot annotations. Alternatively, users have the option to run Liftoff and miniprot separately and then input the annotations using the `‘--liftoff’` and `‘--miniprot’` arguments to LiftOn. The two annotations are then used to create gffutils sqlite3 databases.

### 4.1 Annotation lift-over special handling of the human reference

For all mappings of the human annotation, we excluded all alternative scaffolds and patches from the GRCh38 genome and its annotation. Specifically, we excluded scaffolds ending in “_fix” and “_alt”, because they are duplicates or variants of sequences found on the primary chromosomes. We also excluded rRNAs, which occur in hundreds of identical copies that vary widely among humans and create problems for the alignment programs ^9, 58, 61^.

### 4.2 Pairing Liftoff and miniprot protein-coding transcript annotations

Liftoff was designed to map any genomic interval, containing genes, transcripts, and exons, from one genome onto another. Liftoff’s output uses a hierarchy where genes contain one or more transcripts, and transcripts contain exons and/or CDS features. A CDS feature defines the coding sequences within a protein-coding transcript; therefore, some CDS features span the same intervals as exons, while others span only parts of exons when the exons also include UTRs (untranslated regions). In contrast, miniprot was designed to map protein sequences to a genome and can only generate annotations of transcripts containing CDS features.

After running both Liftoff and miniprot, LiftOn goes through the miniprot mappings and finds, for each one, the Liftoff genomic interval to which it corresponds. A miniprot transcript is considered to match a Liftoff transcript if: (1) the loci of the two transcripts overlap, and (2) the locus of the miniprot transcript must not overlap with any other Liftoff genes loci, except those where the matched Liftoff locus also overlaps. Note that the genomic intervals found by Liftoff are mostly distinct, but it does allow up to 10% overlap between adjacent gene features by default. On the other hand, miniprot lacks the capability to reconcile overlapping gene loci, which sometimes results in a protein-coding transcript from a large gene family to map to multiple gene loci instead of one or to both a pseudogene and the correct locus.

In most cases, LiftOn is able to find a one-to-one mapping between the miniprot transcripts and those from LiftOff. For instance, mapping RefSeq release 220 annotations from GRCh38.p14 to T2T-CHM13 v2 results in 128,351 of 136,978 protein-coding transcripts (93.7%) being uniquely matched between Liftoff and miniprot (See Note S10, and Figure S17).

For the cases where miniprot identifies multiple copies for a protein-coding transcript, LiftOn checks if there at least one copy overlapping with a Liftoff gene locus. Miniprot transcript mappings spanning multiple Liftoff loci are removed to prevent erroneous “read through” annotations. Among the remaining options, the transcript with the highest protein sequence identity score is selected. Note that in the GRCh38.p14 to T2T-CHM13 v2 lift-over there were 355 protein-coding transcripts where none of the miniprot-discovered protein-coding transcripts overlapped with a corresponding Liftoff gene locus.

Once the one-to-one correspondence between two transcripts is established, LiftOn considers both Liftoff and miniprot CDS features of the two transcripts and initiates its protein-maximization algorithm as described in the next Methods section.

### 4.3 Protein-maximization (PM) algorithm

The PM algorithm consists of two components: the chaining algorithm and the open reading frame (ORF) search algorithm. The pseudocode for the `protein_maximization` function is outlined in Algorithm S1.

#### 4.3.1 Step 1: the chaining algorithm

The chaining algorithm is a method for collecting and concatenating the optimal segments of CDSs obtained through the DNA-based mapping approach (Liftoff) and the protein-based approach (miniprot) into a consistent CDS chain, with the goal of maximizing the similarity of the mapped protein to the reference protein. In this context, the reference protein refers to the protein extracted from the reference annotation and genome; e.g., RefSeq annotation on GRCh38.

LiftOn uses Biopython ^59^ to extract and translate mapped protein-coding transcript sequences into their corresponding proteins, and the Parasail ^62^ Python package to align these protein sequences with the reference proteins. Following the alignment step, LiftOn maps the CDS boundaries onto the pairwise protein alignment results (see Figure 1C). The algorithm initially maps the CDS boundaries onto the protein (`get_cds_protein_boundary` function in Algorithm S2) and then adjusts these boundaries using the cigar string obtained from the protein alignment if gaps exist in the target protein (`adjust_cds_protein_boundary` function in Algorithm S2).

Subsequently, the chaining algorithm clusters CDSs from Liftoff and miniprot. It starts by grouping the initial CDSs from both annotations until it encounters a boundary where the aligned amino acid counts in the reference protein up to that boundary match for both Liftoff and miniprot. These CDSs form a comparison block where LiftOn calculates a partial protein sequence identity between the reference protein and those annotated by Liftoff and miniprot, as described in Methods section 4.4. CDS groups with higher identity scores relative to the reference protein are selected for the final LiftOn annotation. After this block is processed, LiftOn identifies a new group of CDSs starting from the previous found boundary and repeats the same selection process. The chaining algorithm’s full pseudocode is provided in Algorithm S3.

#### 4.3.2 Step 2: the open reading frame (ORF) search algorithm

Following the chaining algorithm, LiftOn performs an open reading frame search algorithm on the protein-coding regions of the mapped transcripts that have mutations likely to be more deleterious, such as “frameshift”, “stop codon gain”, “stop codon loss”, and “start codon loss” mutations. The objective is to generate the longest valid protein sequences that align with the full-length reference proteins.

For each selected protein-coding transcript, LiftOn’s open reading frame search algorithm iterates through three reading frames (0, 1, 2). In each frame, it identifies potential ORFs by searching for start codons (“ATG”) and locating the corresponding stop codons (“TAA”, “TAG”, “TGA”), and retains the longest one it finds in that frame.

After identifying potential ORFs, LiftOn proceeds to compare the longest ORF in each frame to the reference protein sequence. It calculates and compares the sequence identity score of each candidate and selects the ORF with the highest sequence identity. If the selected ORF’s identity exceeds the original annotation, LiftOn updates the CDS boundaries of the protein-coding transcript.

### 4.4 DNA and protein transcript sequence identity score calculation

To compute DNA sequence identity scores, LiftOn first extracts transcript sequences by concatenating all exonic regions in a transcript. Subsequently, it aligns each transcript sequence mapped on the target genome by LiftOn, Liftoff, or miniprot to its respective reference sequence. This alignment is performed using the `nw_trace_scan_sat` function from the Parasail ^62^ Python package, configured with a match score of 1, mismatch penalty of −3, gap opening penalty of 2, and gap extension penalty of 2. LiftOn then reports the percent identity between the two aligned sequences, defined as in BLAST ^63^ as the number of matching bases in the two sequences over the number of alignment columns. The pseudocode of the algorithm described in this paragraph is illustrated in the `get_DNA_id_fraction` function from Algorithm S4. An example is provided in Figure S18A.

To compute protein sequence identity scores, LiftOn first generates the protein sequence for each mapped transcript by translating the sequence obtained from concatenating all CDS regions in the transcript. Then, it aligns each protein sequence with the full-length reference protein by performing a pairwise alignment using the BLOSUM 62 matrix ^64^, a gap opening penalty of 11 for insertions and deletions (INDELs), and a gap extension penalty of 2. The protein sequence identity score is calculated up to the first encountered stop codon in the target protein. Differing slightly from the BLAST-style metric employed for DNA sequence identity, LiftOn compresses the gaps in the reference alignment to prevent over-penalization of longer proteins in the target genome. These proteins might be mapped as longer due to potential repeat regions within the target genome or because of potential truncations in the proteins of the reference genome. The pseudocode for this process is presented in the `get_AA_id_fraction` function in Algorithm S4. An example is provided in Figure S18B.

Additionally, the get_partial_id_fraction function in Algorithm S4 describes this process when the sequence identity computation is restricted to evaluating matches between partial proteins, as done in the chaining algorithm, to determine the best matching CDS group to the reference.

### 4.5 Resolving overlapping gene loci and finding extra copy of protein-coding genes

LiftOn employs both Liftoff and miniprot to identify additional copies of protein-coding genes, prioritizing Liftoff for its capability to also map UTR regions. First, LiftOn uses the embedded Liftoff module to iteratively map new gene copies and verify whether the lifted-over gene loci overlap with existing annotations. If so, it allows overlaps that are also present in the reference genome; if not, it permits a maximum overlap of 10% with other gene loci. The Liftoff module also ensures that each gene copy meets the user-specified minimum DNA sequence identity between the reference and target gene loci. This process continues until all possible copies have been identified.

Then, LiftOn applies the miniprot module to identify any additional copies of protein-coding genes that Liftoff might have missed. LiftOn utilizes the intervaltree package, a self-balancing interval tree implemented in Python (https://github.com/chaimleib/intervaltree), to manage gene loci intervals and enable fast detection of overlaps. Since miniprot maps annotations at the protein-coding transcript (mRNA) level, LiftOn copies the corresponding gene-level feature from the reference annotation as the parent of the mRNA feature identified exclusively by miniprot to ensure consistency within the “gene-mRNA-exon” hierarchy. LiftOn also enforces the constraint that any extra gene copies identified by miniprot should have at most a 10% overlap with other gene loci (computed as the ratio of the mapped miniprot length to the smaller of the two values: the mapped miniprot length or the reference protein length).

Furthermore, since miniprot maps only proteins and does not consider the untranslated regions in gene loci, it is very likely that miniprot maps to a processed pseudogene or only maps a partial gene segment. To remove potential pseudogenes, we implemented stricter filtering rules for gene loci identified exclusively by miniprot. First, if miniprot identifies only one CDS (i.e. with no intervening introns), the gene locus is annotated only if there is also a single CDS in the reference annotation. Second, the ratio of the coding regions in the miniprot annotation to the coding regions in the longest isoform of a reference gene locus must be between 0.9 and 1.5. Note that any extra gene copies identified exclusively by miniprot will not include UTRs.

### 4.6 LiftOn arguments used in the study

All analyses in this study were performed using LiftOn with its default parameters along with the additional arguments `‘-copies’`, `‘-sc 0.95’`, and `‘-polish’`. The `‘-copies’` argument prompts Liftoff to search for extra gene copies following the initial lift-over process. A gene copy will be annotated at a specific locus only if there is no overlap with an already annotated feature. Accompanying the `‘-copies’` argument, the `‘-sc 0.95’` parameter modifies the default Liftoff requirement of 100% sequence identity of mapped exons/CDSs to the reference ones, reducing it to allow a 95% sequence identity. The `‘-copies’` argument will also trigger the annotation of extra copies of gene loci, which were distinctly found by miniport.

The `‘-polish’` argument enables the Liftoff step in LiftOn to realign exons to correct CDSs that may have been altered during the lift-over process, such as the loss of start/stop codons or the introduction of in-frame stop codons, although this step extends the processing time.

To map RefSeq release v220 (Results 2.2), MANE version 1.2 (Note S1, Table S4 and Figure S7), and CHESS 3 (Note S2, Table S5 and Figure S8) human annotations from GRCh38 to CHM13 (Table 1 and Figure 2), the `‘-chroms <chromosome_mapping.txt>’` argument was also employed in order to first perform a mapping step on a chromosome-by-chromosome basis. Once this step is complete, any genes that remain unmapped are remapped to the entire target assembly. This setting resulted in improved human annotation mapping accuracy. For all other experiments, LiftOn was run without the `‘-chroms’` argument, mapping gene sequences to the full genome.

The `‘-f’` argument and the input file `‘features.txt’` indicates all types of parent features to lift over. To ensure LiftOn does not mistakenly identify pseudogenes as extra gene copies, we ran LiftOn with the `‘features.txt’` file specifying both “gene” and “pseudogene” features, to lift them over along with their children.

LiftOn runs miniprot with default parameters plus the `‘-gff-only’` option to exclusively generate a GFF output file.

All experiments were conducted on a 24-core, 48-thread Intel(R) Xeon(R) Gold 6248R Linux computer with 1024 GB memory, using a single thread of execution.

### 4.7 Gene and transcript feature counting

The counts of protein-coding and non-coding genes and transcripts reported in this study were calculated as follows:

- Gene counting Gene features were classified as “protein-coding”, “non-coding”, and “others” based on the “gene_biotype” attribute in NCBI’s RefSeq or the “gene_type” attribute in EMBL-EBI’s Ensembl/GENCODE and CHESS 3. More specifically, a gene feature was categorized as a protein-coding gene if the feature type was “gene” and its type attribute was “protein_coding”; a gene was categorized as non-coding if its type attribute was either “lncRNA” or “ncRNA”. Other gene features, including “Pseudogene”, “miRNA”, “snoRNA”, “tRNA”, “V_segment”, “snRNA”, “J_segment”, “misc_RNA”, “C_region”, “antisense_RNA”, etc., were categorized as “others”. The full list is provided in Table S1.
- Transcript counting Transcript features were also categorized into “protein-coding”, “non-coding”, and “others”. The criteria for classification were based on both the type of transcript and the type of its parent gene. A transcript was counted as protein-coding if its feature type was “mRNA”, and its parent gene was a protein-coding gene; a transcript was classified as non-coding if its feature type was either “lncRNA” or “ncRNA” and its parent gene was also a non-coding gene. The remaining transcripts were categorized as “others”.

### 4.8 Running LiftOn on a test dataset

We provide sample files for the users wanting to test LiftOn at our GitHub repository: https://github.com/Kuanhao-Chao/LiftOn/tree/main/test. To execute LiftOn, the user can type a single command that uses the `‘-g’` flag to specify a reference annotation file in GFF format (e.g.,

`‘NCBI_RefSeq_no_rRNA.gff’`), the `‘-O’` flag to specify the output file (e.g.,

`‘lifton.gff3’`), plus both the reference genome file (e.g.,

`‘GCF_000001405.40_GRCh38.p14_genomic.fn’`) and the target genome file (e.g.,

`‘chm13v2.0.f’`). e.g.:

`lifton -g NCBI_RefSeq_no_rRNA.gff -o lifton.gff3 -copies chm13v2.0.fa GCF_000001405.40_GRCh38.p14_genomic.fna`

### 4.9 Genomes used in this study

Brief rationale for selecting the genomes used in our experiments are described below.

#### Apis mellifera

At the time of this study, GenBank contained no *Apis mellifera* assemblies of the same strain as the reference genome, Amel_HAv3.1 (Strain DH4) ^40^. We considered both ASM1932182v1 ^39^ (strain *ligustica*) and ASM1384124v2 (strain *carnica*), and chose ASM1932182v1 because it was more contiguous (42 contigs) than ASM1384124v2 (313 contigs).

#### Arabidopsis thaliana

The *A. thaliana* reference genome, TAIR10.1 ^43^, represents the Columbia (Col-0) strain. Of the other Col-0 assemblies, Col-CC and Col-CEN (ASM2311539v1) ^44^ were of similar quality, having all chromosomes completely assembled. We chose Col-CEN (ASM2311539v1) because Col-CC was a consensus assembly of 13 independent assemblies, whereas Col-CEN (ASM2311539v1) was derived from a single organism.

#### Mus musculus

At the time of this study, GenBank contained no high-quality contiguous assemblies of the same strain (C57BL/6J) as the reference genome, GRCm39 ^56^. The only other assembly of the C57BL/6J strain (ASM377452v2) is quite fragmented, with 36,193 contigs. We chose NOD_SCID ^37^ (strain NOD/SCID) instead because it has the fewest contigs (281) of all other *M. musculus* assemblies from known strains.

#### Oryza sativa

The reference genome for rice, IRGSP-1.0 ^42^, belongs to the Japonica subgroup and the Nipponbare cultivar. From the three other assemblies that were also from the Nipponbare cultivar, we chose ASM3414082v1 ^41^ as it was the most contiguous, with every chromosome fully assembled into a single scaffold.

## 5. Data and Code availability

LiftOn is implemented as a Python package. The LiftOn project is freely available on github from: github.com/Kuanhao-Chao/LiftOn, and on PyPi from: https://pypi.org/project/LiftOn/. The LiftOn documentation is available at: ccb.jhu.edu/lifton.

## 6. Author Contribution

K.C., M.P., and S.L.S. designed the research.

K.C. and A.S. developed the LiftOn software. K.C., J.H., A.M., M.P., and S.L.S. analyzed the results.

K.C. and J.H. conducted experiments on Liftoff and miniprot annotation mapping, involving both the same species and species that were closely and distantly related.

K.C. and C.H. visualized the results.

K.C. and A.M. wrote the LiftOn documentation.

J.H. devised the name “LiftOn” for the software. A.M. designed the LiftOn logo.

K.C., J.H., C.H., M.P., and S.L.S. wrote the manuscript.

## Supporting information

Supplementary Algorithms

