## Supplementary Notes for "Combining DNA and protein alignments to improve genome annotation with LiftOn"

### **Note S1.** Mapping RefSeq MANE v1.2 annotation from GRCh38 to T2T-CHM13

We mapped RefSeq MANE[1] version 1.2 of GRCh38 annotation from GRCh38.p14 onto T2T-CHM13 v2.0. To accurately report the results from LiftOn's open-reading frame search module, we excluded specific exceptions, such as alternate start codons and selenoproteins, during the lift-over process. We also excluded genes on the fixed and genome patches in the GRCh38 annotation file, specifically those with the UCSC naming style ending in "_fix" and "_alt", as they are duplicates of genes found on the primary chromosomes. The genomic distance between the two genomes is 0.93 Dashing[2] similarity score. The distance estimated by Mash[3] is 0.000966693.

The command to map “gene” and “pseudogene” features in the MANE annotation from GRCh38.p14 onto T2T-CHM13 v2.0, chromosome by chromosome, is provided below. For a more detailed explanation of the arguments, please refer to the documentation at: https://ccb.jhu.edu/lifton/.

```
lifton -g MANE.GRCh38.v1.2.refseq_genomic.cleaned_no_alt_no_fix.gff -o
/output/path/lifton.gff3 -polish -copies -sc 0.95 chm13v2.0.fa
GCF_000001405.40_GRCh38.p14_genomic.fna -chroms
human_chroms_mapping.csv -f features.txt
```

Figure S7A illustrates the genome annotation discrepancies between Liftoff and miniprot, while LiftOn shows improvements over both Liftoff (as depicted in Figure S7B) and miniprot (as shown in Figure S7C). Frequency plots for the three tools are presented from left to right in Figure S7D. LiftOn identified 24 proteins with a protein sequence identity score below 0.4, in contrast to Liftoff's 156 and miniprot’s 467.

LiftOn successfully mapped 19,130 genes onto CHM13, achieving a 99.9% mapping rate, with 28 genes remaining unmapped. LiftOn identified 162 genes with additional copies (See Table S4 and Figures S7E,F). In total, LiftOn mapped 19,535 gene loci from GRCh38 to T2T-CHM13. The protein-coding gene order plot is shown in Figure S7G.

**Note S2.** Mapping CHESS 3 annotation from GRCh38 to T2T-CHM13*:*

We mapped CHESS 3 [4] of GRCh38 annotation (available at https://ccb.jhu.edu/chess/) from GRCh38.p12 onto T2T-CHM13 v2.0. Since the CHESS 3 annotation hierarchy does not have a gene level, all features are structured as “transcript – exon / CDS”. Therefore, LiftOn conducted the lift-over at the transcript level.

Following the same approach as MANE, we also excluded protein-coding transcripts on the genome patches in the GRCh38 annotation file, specifically those with the UCSC naming style ending in "_alt", as they are duplicates of genes found on the primary chromosomes.

The command to map “transcript” features in the CHESS 3 annotation from GRCh38.p14 onto T2T-CHM13 v2.0, chromosome by chromosome, is provided below. For a more detailed explanation of the arguments, please refer to the documentation at: https://ccb.jhu.edu/lifton/.

```
lifton -g chess3.0.1.gff -o /output/path/lifton.gff3 -polish -copies -
sc 0.95 chm13v2.0.fa hg38_p12_ucsc.no_alts.no_fixs.fa -chroms
human_chroms_mapping.csv -f features.txt
```

LiftOn successfully mapped 99,506 protein-coding transcripts onto CHM13, achieving a 94.5% mapping rate. The results of the LiftOn CHESS 3 transcript mapping are summarized in Table S5 and Figure S8.

**Note S3.** Mapping genes between two *Mus musculus* (house mouse) genomes

We mapped RefSeq GRCm39 annotation (GCF_000001635.27-RS_2023_04) from GRCm39 [5] (GCF_000001635.27l, C57BL/6J strain, https://www.ncbi.nlm.nih.gov/datasets/genome/GCF_000001635.27/) onto NOD_SCID [6] assembly (GCA_031763685.1, NOD/SCID strain, https://www.ncbi.nlm.nih.gov/datasets/genome/GCA_031763685.1/). The genomic distance between the two genomes is 0.86 Dashing [2] similarity score. The distance estimated by Mash [3] is 0.00245807.

The command to map "gene" and "pseudogene" features in RefSeq GRCm39 annotation from GRCm39 onto NOD_SCID assembly is provided below. For a more detailed explanation of the arguments, please refer to the documentation at: https://ccb.jhu.edu/lifton/.

```
lifton -g GRCm39_genomic.gff -o /output/path/lifton.gff3 -polish -
copies -sc 0.95 NOD_SCID_genomic.fna GRCm39_genomic.fna -f
features.txt
```

At the gene level, 35,031 genes were successfully mapped, with 520 genes remaining unmapped, including 487 protein-coding and 33 non-coding genes. The total gene mapping rate is 98.5%.

At the transcript level, 118,906 transcripts were successfully mapped, and 839 transcripts were unmapped, comprising 743 protein-coding and 96 non-coding transcripts. The overall transcript mapping rate is also 99.3%. The results of the LiftOn annotation mapping between two mouse genomes are summarized in Table S6 and illustrated in Figure S9.

### **Note S4.** Mapping between two *Apis mellifera* (honey bee) genomes

We mapped NCBI RefSeq *Apis mellifera* annotation release 104 from Amel_HAv3.1 [7] (GCF_003254395.2, DH4 strain. https://www.ncbi.nlm.nih.gov/datasets/genome/GCF_003254395.2/) onto ASM1932182v1 assembly [8] (GCA_019321825.1, ligustica strain. https://www.ncbi.nlm.nih.gov/datasets/genome/GCA_019321825.1/). The genomic distance between the two genomes is 0.75 Dashing [2] similarity score. The distance estimated by Mash [3] is 0.00484456.

The command to map "gene" and "pseudogene" features in *Apis mellifera* annotation release 104 from Amel_HAv3.1 onto ASM1932182v1 assembly is provided below. For a more detailed explanation of the arguments, please refer to the documentation at: https://ccb.jhu.edu/lifton/.

```
lifton -g HAv3.1_genomic.gff -o /output/path/lifton.gff3 -polish -
copies -sc 0.95 ASM1932182v1_genomic.fna Amel_HAv3.1_genomic.fna -f
features.txt
```

At the gene level, 11,740 genes were successfully mapped, with 53 genes remaining unmapped, including 40 protein-coding and 13 non-coding genes. The gene mapping rate is 99.6%.

At the transcript level, 26,493 transcripts were successfully mapped, and 124 transcripts were unmapped, comprising 101 protein-coding and 23 non-coding transcripts. The overall transcript mapping rate is also 99.5%. The results of the LiftOn annotation mapping between two honey bee genomes are summarized in Table S7 and illustrated in Figure S10.

### **Note S5.** Mapping genes between two *Oryza sativa* (Asian rice) genomes

We mapped NCBI RefSeq *Oryza sativa* Japonica Group annotation release 102 from IRGSP-1.0 [9] (GCF_001433935.1, https://www.ncbi.nlm.nih.gov/datasets/genome/GCF_001433935.1/) onto ASM3414082v1 assembly [10] (GCA_034140825.1, https://www.ncbi.nlm.nih.gov/datasets/genome/GCA_034140825.1/). The genomic distance between the two genomes is 0.99 Dashing [2] similarity score. The distance estimated by Mash [3] is 0.000177174.

The command to map "gene" and "pseudogene" features in RefSeq *Oryza sativa* Japonica Group annotation release 102 from IRGSP-1.0 onto ASM3414082v1 assembly is provided below. For a more detailed explanation of the arguments, please refer to the documentation at: https://ccb.jhu.edu/lifton/.

```
lifton -g IRGSP_genomic.gff -o /output/path/lifton.gff3 -polish -
copies -sc 0.95 ASM3414082v1_genomic.fna IRGSP_genomic.fna -f
features.txt
```

At the gene level, 32,008 genes were successfully mapped, with 36 genes remaining unmapped, including 31 protein-coding and 5 non-coding genes. The total gene mapping rate is 99.9%.

At the transcript level, 48,843 transcripts were successfully mapped, and 45 transcripts were unmapped, comprising 39 protein-coding and 6 non-coding transcripts. The overall transcript mapping rate is also 99.9%. The results of the LiftOn annotation mapping between two rice genomes are summarized in Table S8 and illustrated in Figure S11.

**Note S6.** Mapping genes between two *Arabidopsis thaliana* (thale cress) genomes

We mapped RefSeq TAIR10.1 annotation (submitted by TAIR and Araport) from TAIR10.1 [11] (GCF_000001735.4, https://www.ncbi.nlm.nih.gov/datasets/genome/GCF_000001735.4/) onto ASM2311539v1 assembly [12] (GCA_023115395.1, https://www.ncbi.nlm.nih.gov/datasets/genome/GCA_023115395.1/). The genomic distance between the two genomes is 0.99 Dashing [2] similarity score. The distance estimated by Mash [3] is 0.000133896.

The command to map "gene" and "pseudogene" features in RefSeq TAIR10.1 annotation from TAIR10.1 onto ASM2311539v1 assembly is provided below. For a more detailed explanation of the arguments, please refer to the documentation at: https://ccb.jhu.edu/lifton/.

```
lifton -g TAIR10.gff -o /output/path/lifton.gff3 -polish -copies -sc
0.95 ASM2311539v1_genomic.fna TAIR10.fna -f features.txt
```

At the gene level, 31,227 genes were successfully mapped, with 254 genes remaining unmapped, including 38 protein-coding and 216 non-coding genes. The total gene mapping rate is 99.2%.

At the transcript level, 52,369 transcripts were successfully mapped, and 266 transcripts were unmapped, comprising 50 protein-coding and 216 non-coding transcripts. The overall transcript mapping rate is also 99.5%. The results of the LiftOn annotation mapping between two thale cress genomes are summarized in Table S9 and illustrated in Figure S12.

**Note S7.** Mapping RefSeq annotations from *Homo sapiens* genome (GRCh38) to *Pan troglodytes* genome (chimpanzee)

We chose NCBI RefSeq [13] release v220 of GRCh38.p14, excluding rRNAs [14], the same version used in the GRCh38-to-CHM13 lift-over experiment. This was used for mapping annotations from GRCh38 onto NHGRI_mPanTro3-v1.1-hic.freeze_pri (GCF_028858775.1, https://www.ncbi.nlm.nih.gov/datasets/genome/GCF_028858775.1/). The genomic distance between the two genomes is 0.47 Dashing [2] similarity score. The distance estimated by Mash [3] is 0.0130253.

The command to map "gene" and "pseudogene" features in RefSeq release 220 annotations from GRCh38 onto NHGRI_mPanTro3-v1.1-hic.freeze_pri is provided below. For a more detailed explanation of the arguments, please refer to the documentation at: https://ccb.jhu.edu/lifton/.

```
lifton -g NCBI_RefSeq_no_rRNA_no_alt_no_fix.gff -o
/output/path/lifton.gff3 -polish -copies -sc 0.95 NHGRI_mPanTro3-
v1.1.fna GCF_000001405.40_GRCh38.p14_genomic.fna -f features.txt
```

At the gene level, 37,509 genes were successfully mapped, with 477 genes remaining unmapped, including 285 protein-coding and 192 non-coding genes. The total gene mapping rate is 98.7%.

At the transcript level, 160,561 transcripts were successfully mapped, and 1,615 transcripts were unmapped, comprising 1,219 protein-coding and 396 non-coding transcripts. The overall transcript mapping rate is also 99.0%. The results of the LiftOn annotation mapping between human and chimpanzee genomes are illustrated in Figure S13.

**Note S8.** Mapping from *Drosophila melanogaster* to *Drosophila erecta*

We mapped the *D .melanogaster* RefSeq FlyBase Release 6.54 annotation from RefSeq release 6 plus ISO1 MT assembly [15] (GCF_000001215.4, dm6 strain, https://www.ncbi.nlm.nih.gov/datasets/genome/GCF_000001215.4/) onto the *D. erecta* DereRS2 assembly (GCA_003286155.2, 14021-0224.00,06,07 strain, https://www.ncbi.nlm.nih.gov/datasets/genome/GCA_003286155.2/). The genomic distance between the two genomes is 0.07 Dashing [2] similarity score. The distance estimated by Mash [3] is 0.0769152.

The command to map "gene" and "pseudogene" features in FlyBase Release 6.54 annotation from RefSeq release 6 plus ISO1 MT assembly onto the *D. erecta* DereRS2 assembly is provided below. For a more detailed explanation of the arguments, please refer to the documentation at: https://ccb.jhu.edu/lifton/.

```
lifton -g d.menogaster_genomic.gff -o /output/path/lifton.gff3 -polish
-copies -sc 0.95 d.erecta_genomic_bk.fna d.menogaster_genomic.fna -f
features.txt
```

At the gene level, 15,124 genes were successfully mapped, with 881 genes remaining unmapped, including 642 protein-coding and 239 non-coding genes. The total gene mapping rate is 94.5%.

At the transcript level, 31,993 transcripts were successfully mapped, and 1,183 transcripts were unmapped, comprising 872 protein-coding and 311 non-coding transcripts. The overall transcript mapping rate is also 96.4%. The results of the LiftOn annotation mapping between *Drosophila m.* and *Drosophila e.* genomes are illustrated in Figure S14.

**Note S9.** Mapping from *Mus musculus* (GRCm39) to *Rattus norvegicus* (mRatBN7.2)

We mapped RefSeq GRCm39 annotation (GCF_000001635.27-RS_2023_04) from GRCm39 [5] (GCF_000001635.27l, C57BL/6J strain. https://www.ncbi.nlm.nih.gov/datasets/genome/GCF_000001635.27/) onto mRatBN7.2 [16] (GCF_015227675.2, BN/NHsdMcwistrain, https://www.ncbi.nlm.nih.gov/datasets/genome/GCF_015227675.2/). The genomic distance between the two genomes is 0.01 Dashing [2] similarity score. The distance estimated by Mash [3] is 0.120127.

The command to map "gene" and "pseudogene" features in RefSeq GRCm39 annotation from GRCm39 onto mRatBN7.2 is provided below. For a more detailed explanation of the arguments, please refer to the documentation at: https://ccb.jhu.edu/lifton/.

```
lifton -g IRGSP_genomic.gff -o /output/path/lifton.gff3 -polish -
copies -sc 0.95 ASM3414082v1_genomic.fna IRGSP_genomic.fna -f
features.txt
```

At the gene level, 33,542 genes were successfully mapped, with 2,009 genes remaining unmapped, including 1,576 protein-coding and 433 non-coding genes. The total gene mapping rate is 94.3%.

At the transcript level, 115,692 transcripts were successfully mapped, and 4,053 transcripts were unmapped, comprising 3,300 protein-coding and 753 non-coding transcripts. The overall transcript mapping rate is also 96.6%. The results of the LiftOn annotation mapping between *Drosophila m.* and *Drosophila e.* genomes are illustrated in Figure S15.

### **Note S10.** Analysis of gene loci consensus between Liftoff and miniprot of mapping RefSeq release 220 annotation from GRCh38 to T2T-CHM13

In this analysis, we have two goals: (1) to determine how many protein-coding genes miniprot can map that are missed by Liftoff, and (2) to assess the degree of consensus between Liftoff and miniprot regarding the coordinates of protein-coding gene loci. We investigated the results obtained by mapping the RefSeq release 220 annotations from GRCh38.p14 to T2T-CHM13 v2.

We ran Liftoff with the additional arguments '-copies', '-sc 0.95', and '-polish', and specified '-chroms <chromosome_mapping.txt>' to conducted chromosome-to-chromosome gene loci mapping. We ran miniprot with default parameters and with the argument '-gff-only' to generate a GFF file only.

Liftoff maintains the same gene/transcript IDs as those of the mapped gene loci, whereas miniprot assigns a new "MP<transcript_number>" as the transcript ID, and original gene/transcript IDs as as the "Target" attribute value. To pair genes in these two annotations, we created dictionary mappings from reference IDs to miniprot IDs. Subsequently, we used gffutils to generate sqlite3 databases for both Liftoff and miniprot annotations.

We began by iterating through the protein IDs mapped by miniprot to identify corresponding Liftoff transcripts and any all copies identified by miniprot.

Mappings were categorized as follows: "one-to-one mapping" when a single miniprot transcript overlapped with a Liftoff transcript; "one-to-many mapping" when multiple miniprot transcripts existed and at least one overlapped with a Liftoff transcript; "Liftoff- miniprot disagreement" when one or more miniprot transcripts were recognized but none overlapped with a Liftoff transcript; "Liftoff misses" when miniprot identified transcripts not detected by Liftoff.

The majority of cases are "1-to-1 mapping", comprising a total of 128,351 protein-coding transcripts. 1,986 protein-coding transcript loci are categorized under "one-to-many mapping", with miniprot identifying a total of 7,150 transcripts. Additionally, there are 355 protein-coding transcript loci that fall under "Liftoff-miniprot disagreement" and 334 loci under "Liftoff misses".

For protein-coding transcripts in the "1-to-1 mapping" category, LiftOn can directly pair them and run the protein-maximization algorithm. For protein-coding transcripts in the "one-to-many mapping" category, LiftOn needs to first find the most overlapping one, pair them, and run the protein-maximization algorithm. When multiple miniprot transcripts overlap with a Liftoff transcript, we initially check if any of them span more than one distinct locus. Any mapped transcripts that cross multiple gene loci are removed to eliminate miniprot-generated "read through" annotations. If multiple miniprot transcripts still remain, the one with the higher protein sequence identity score is selected. The remaining extra protein-coding transcripts in the "Liftoff-miniprot disagreement" and "Liftoff misses" categories are used to identify additional copies.

In summary, Figure S17A illustrates the alignment of 130,337 (128,351+1,986) miniprot-identified protein-coding gene loci with their respective Liftoff gene loci. Conversely, Figure S17B depicts 5,925 protein-coding transcripts within 1,986 miniprot-identified loci, which are unique copies exclusive to miniprot and do not overlap with Liftoff loci.
