## Supplementary Figures for "Combining DNA and protein alignments to improve genome annotation with LiftOn"

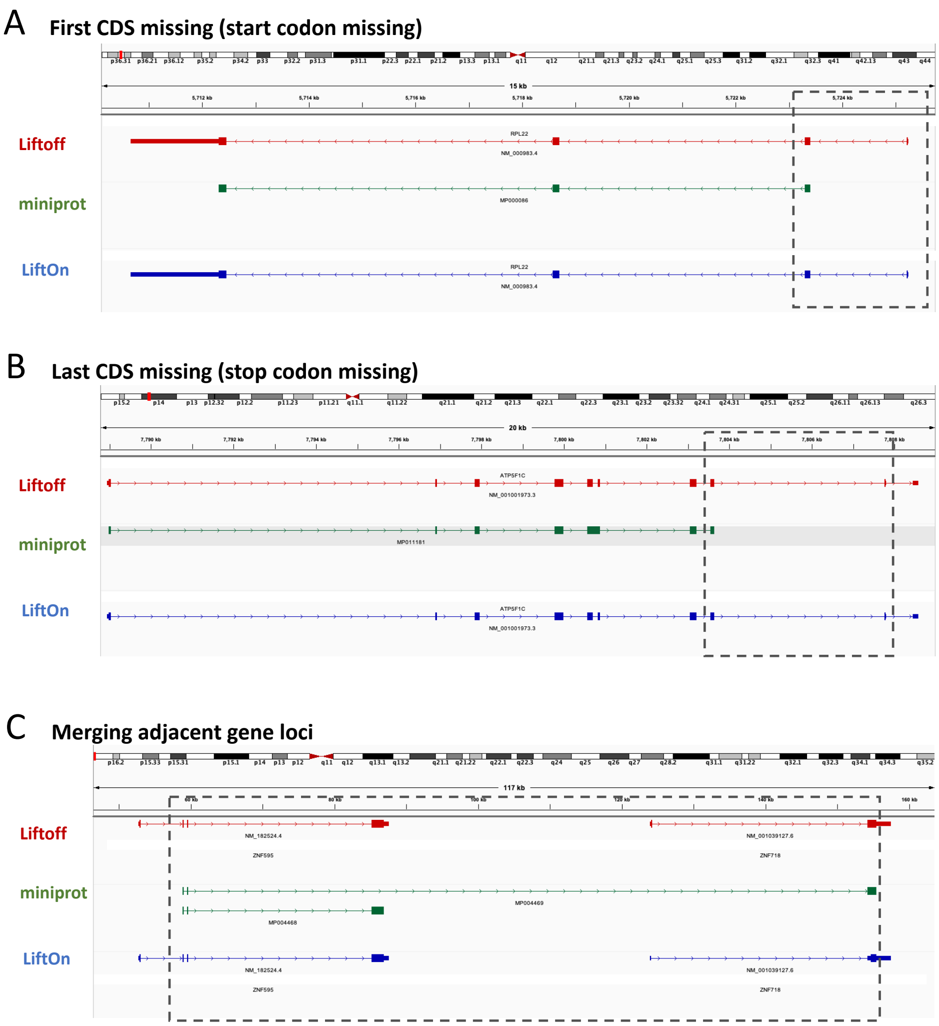


RPL22 (NM_000983.4) / MP000086
chr1:5710666-5725241

| | Protein identity |
|---|---|
| Liftoff | 100% |
| miniprot | 96.90% |
| LiftOn | 100% |

ATP5F1C (NM_001001973.3) / MP011181
chr10:7788947-7808583

| | Protein identity |
|---|---|
| Liftoff | 76.70% |
| miniprot | 99.55% |
| LiftOn | 99.55% |

ZNF718 (NM_001039127.6) / MP004468
chr4:123941-157507

| | Protein identity |
|---|---|
| Liftoff | 0.60% |
| miniprot | 93.80% |
| LiftOn | 12.47% |

ZNF595 (NM_182524.4) / MP004469
chr4:52753-87637

| | Protein identity |
|---|---|
| Liftoff | 100% |
| miniprot | 99.85% |
| LiftOn | 100% |

**Figure S1.** Common mistakes found in miniprot alignment **(A)** miniprot exhibits a tendency to miss the first overhanging small coding sequences (CDS), leading to start codon missing mutations. This is highlighted in the dashed box for the NM_000983.4 transcript in the RPL22 gene locus (chr1:5710666-5725241). Protein sequence identity scores for Liftoff, miniprot, and LiftOn are recorded at 100%, 96.90%, and 100%, respectively. **(B)** miniprot exhibits a tendency to miss the last CDS, resulting in stop codon missing or premature stop codon mutations. This is exemplified in the dashed box representing the NM_001001973.3 transcript in the ATP5F1C gene locus (chr1:7788947-7808583). The corresponding protein sequence identity scores for Liftoff, miniprot, and LiftOn are documented as 76.70%, 99.50%, and 99.50%, respectively. **(C)** For genes in the same gene family adjacent to each other, miniprot may erroneously amalgamate CDSs. The highlighted dashed box exemplifies the miniprot transcript MP011181, linking NM_182524.4 in the ZNF595 gene locus (chr4:123941-157507) and NM_001039127.6 in the ZNF718 gene locus (chr4:52753-87637), resulting in a false run-through transcript annotation.

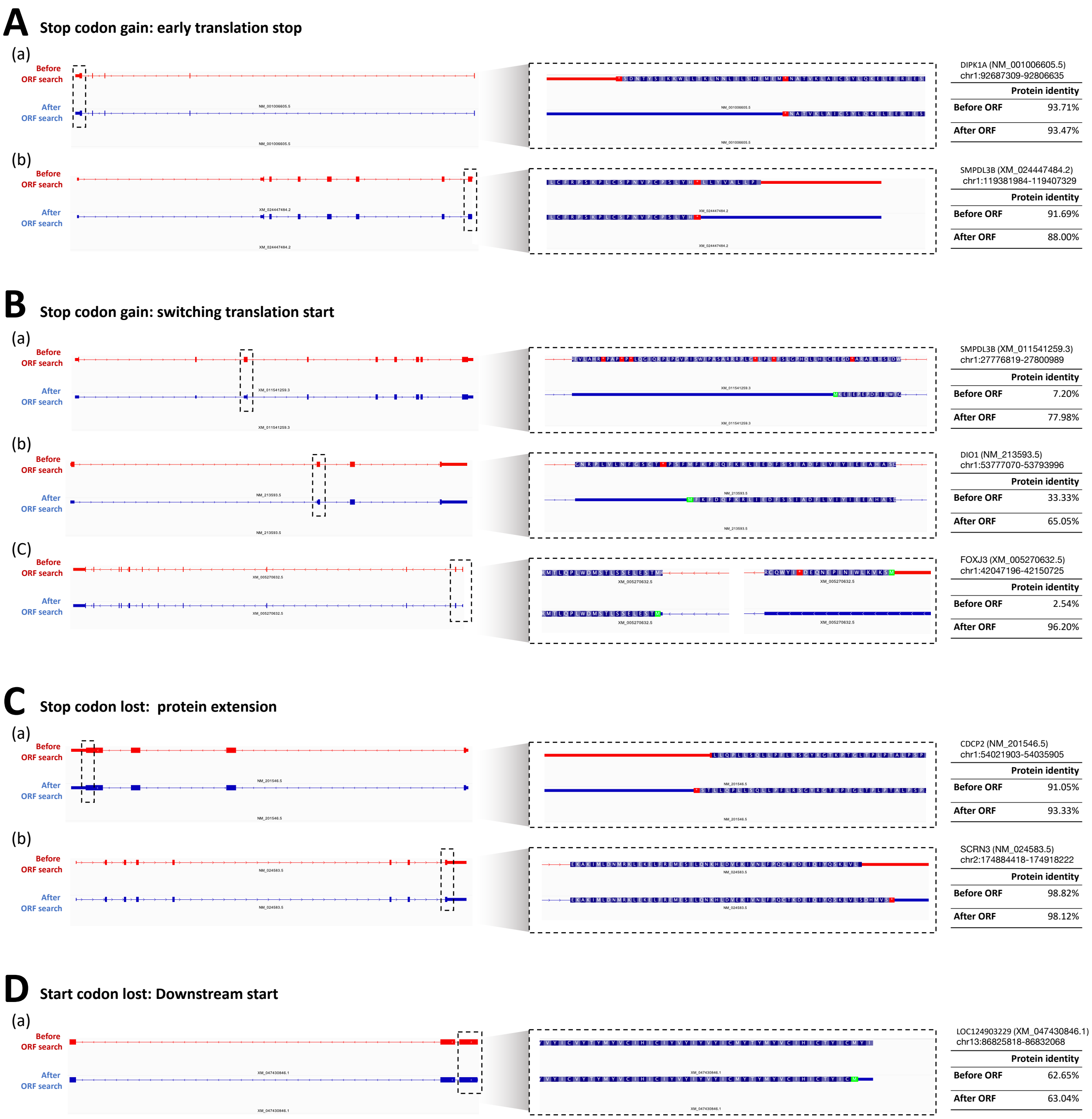


**Figure S2.** Examples of transcripts illustrating different results before and after LiftOn's *open-reading frame search algorithm*. **(A)** Two instances of stop codon gain leading to early translation termination are presented. **(A.a)** In the NM_001006605.5 transcript of the DIPK1A gene (chr1:92687309-92806635), a premature stop codon is identified in the last coding sequence (CDS) in addition to the original stop codon. LiftOn adjusts the CDS end boundary to the upstream first-encountered stop codon. Protein sequence identities before and after ORF search are 93.7% and 93.5%, respectively. **(A.b)** In the XM_024447484.2 transcript of the SMPDL3B gene (chr1:119381984-119407329), the originally annotated transcript lacks a stop codon, and a stop codon is found 26 nucleotides upstream. Protein sequence identities before and after ORF search are 91.7% and 88.0%, respectively. **(B)** Three instances of stop codon gain. By switching to the downstream translation start, the transcript can retain the longer protein length compared to its corresponding protein. **(B.a)** The XM_011541259.3 transcript of the SMPDL3B gene (chr1:27776819-27800989). Protein sequence identities before and after ORF search are 7.2% and 80.0%, respectively. **(B.b)** The NM_213593.5 transcript of the DIO1 gene (chr1:53777070-53793996). Protein sequence identities before and after ORF search are 33.3% and 65.1%, respectively. **(B.c)** The XM_005270632.5 transcript of the FOXJ3 gene (chr1:42047196-42150725). Protein sequence identities before and after ORF search are 2.5% and 96.2%, respectively. **(C)** Two instances of stop codon loss. The ORF search algorithm extends the original protein annotation by expanding the coding sequence (CDS) end boundary to the 3' downstream stop codon. **(C.a)** The NM_201546.5 transcript of the CDCP2 gene (chr1:54021903-54035905). The ORF search algorithm extends the original protein by two amino acids and introduces a stop codon. Protein sequence identities before and after ORF search are 91.1% and 93.3%, respectively. **(C.b)** The NM_024583.5 transcript of the SCRN3 gene (chr2:174884418-174918222). The ORF search extends the original protein by six amino acids and introduces a stop codon. Protein sequence identities before and after ORF search are 98.8% and 98.1%, respectively. **(D)** The XM_047430846.1 transcript of the LOC124903229 gene (chr13:86825818-86832068) is an example of a missing start codon. The ORF search algorithm identifies a valid start codon two amino acids downstream.

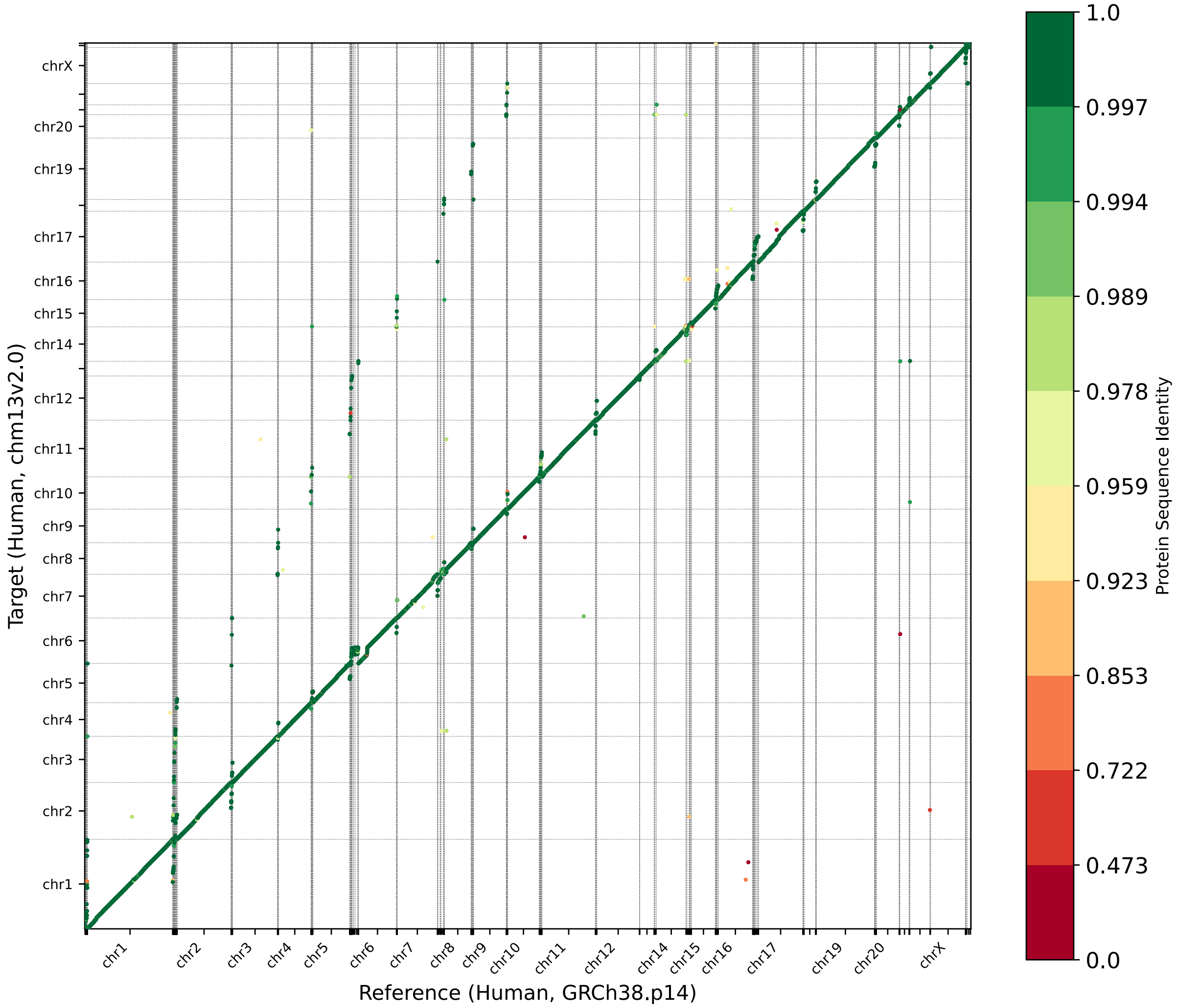


**Figure S3.** Protein-gene order plot, with the x-axis representing the reference genome (GRCh38) and the y-axis representing the target genome (T2T-CHM13). The protein sequence identities are color-coded on a logarithmic scale, ranging from green to red. Green represents a sequence identity score of 1, while red corresponds to a sequence identity score of 0.

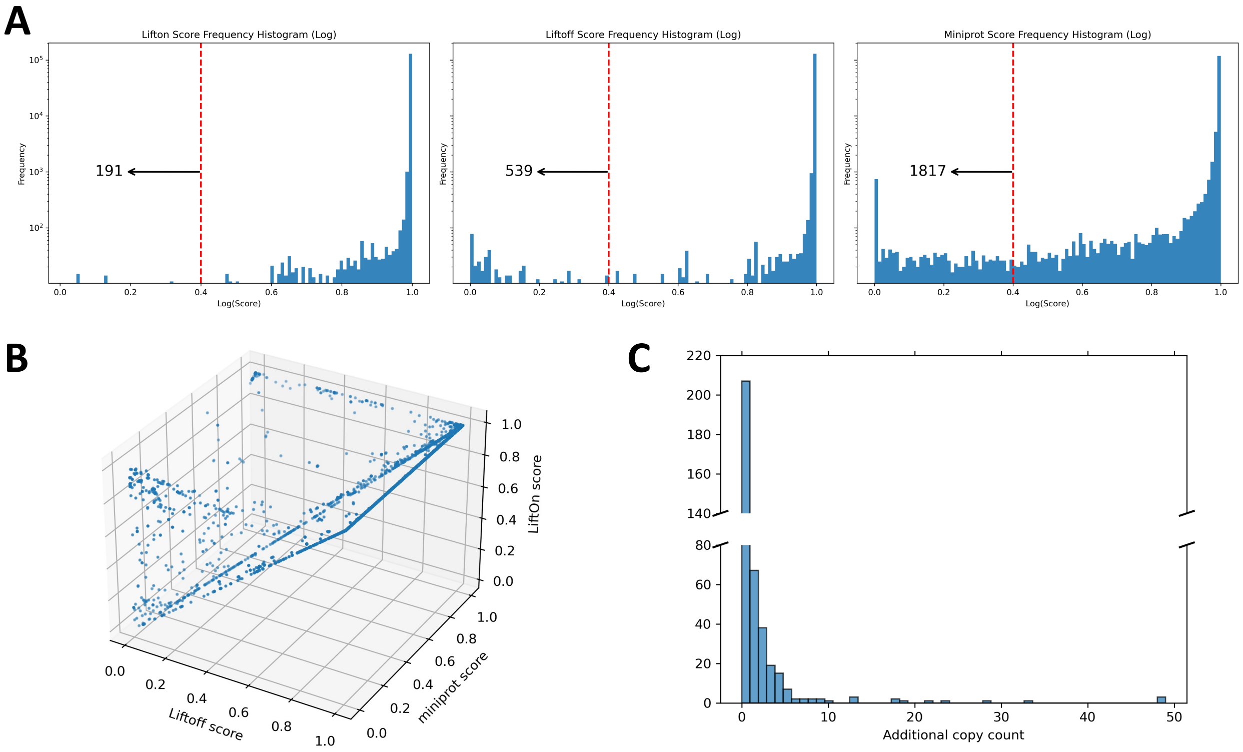


**Figure S4.** Comparative analysis of mapping RefSeq release 220 annotation from GRCh38.p14 to T2T-CHM13 v2.0. **(A)** Frequency plot in logarithmic scale of protein sequence identity for LiftOn (left), Liftoff (middle), and miniprot (right). The number of truncated genes with a protein sequence identity lower than 0.4 threshold, as indicated by the red dotted line, is 191 for LiftOn, 539 for Liftoff, and 1817 for miniprot. **(B)** 3-D scatter plot comparing Liftoff (x-axis), miniprot (y-axis), and LiftOn (z-axis). Each dot represents a protein-coding transcript. If it is above the $x = y$ plane, it indicates that the LiftOn annotation possesses a higher protein sequence identity score and corresponds to a longer protein that aligns with the proteins in the reference annotation. **(C)** Frequency plot of extra gene copy counts reported by LiftOn.

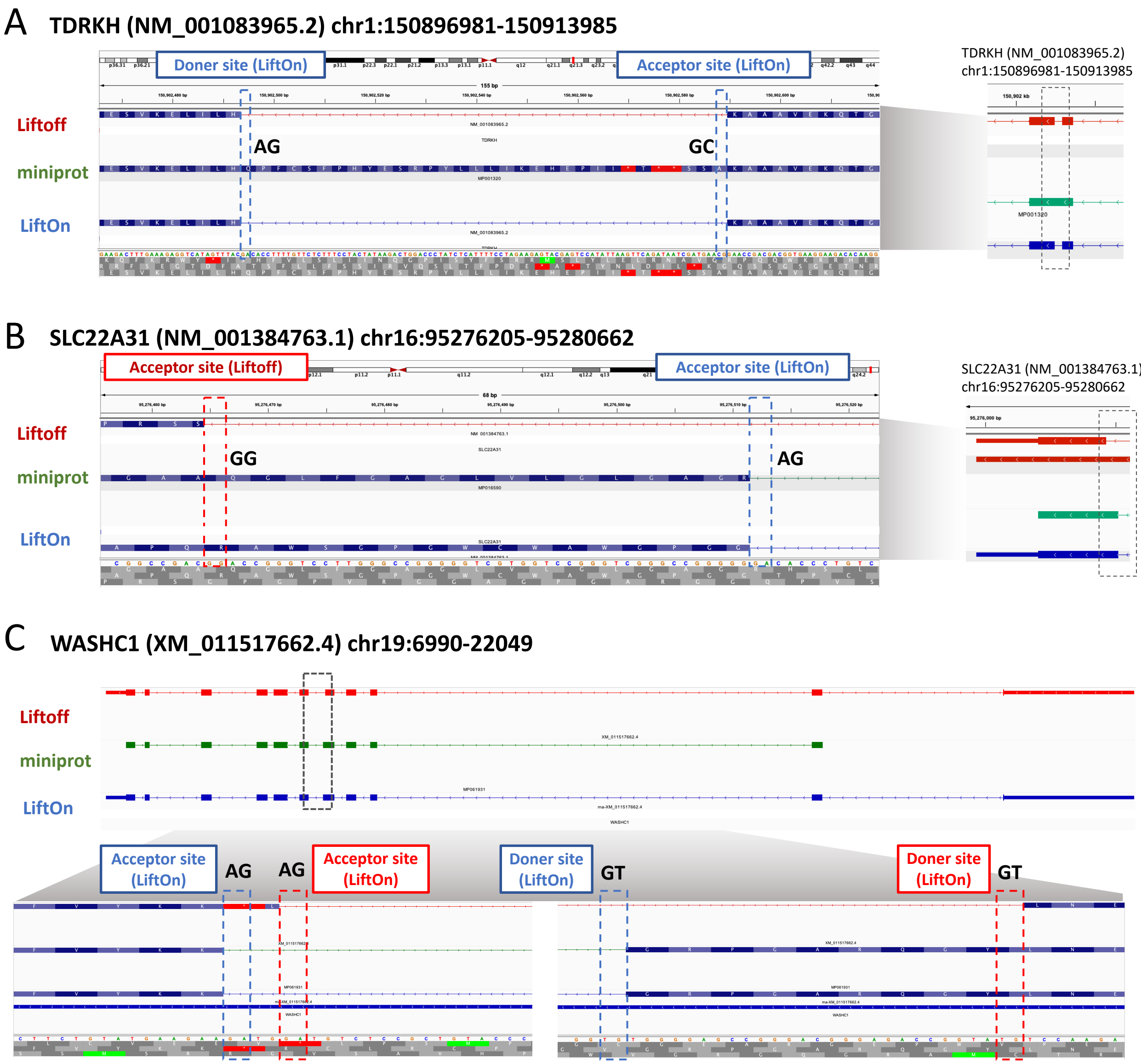


**Figure S5.** The zoomed-in IGV screenshots of Figure 2 depict the following details. **(A)** The splice junction in LiftOn and Liftoff annotations exhibits a donor motif of "GC" and an acceptor motif of "AG". The retained intron in the miniprot annotation introduces a premature stop codon. **(B)** There is a discrepancy between Liftoff and miniprot regarding the acceptor site. Liftoff identifies the acceptor site motif as "GG," while miniprot identifies it as "AG." In this example, LiftOn adopts the splice site information from miniprot. **(C)** In transcript XM_011517662.4 in the WASHC1 gene (chr19:6990 - 22049), the highlighted splice junctions in both Liftoff and miniprot preserve the motif "GT" for donor sites and "AG" for acceptor sites, representing canonical splice sites. However, the splice junction in Liftoff introduces a premature stop codon in the fifth coding sequence (CDS), resulting in a truncated protein with 38.9% protein sequence identity. On the other hand, miniprot preserves 99.14% of the protein but lacks the first overhanging small CDS. Consequently, for the discordant splice junction, LiftOn adopts the one from and rectifies the absence of the first CDS by incorporating Liftoff's CDS.

**A**

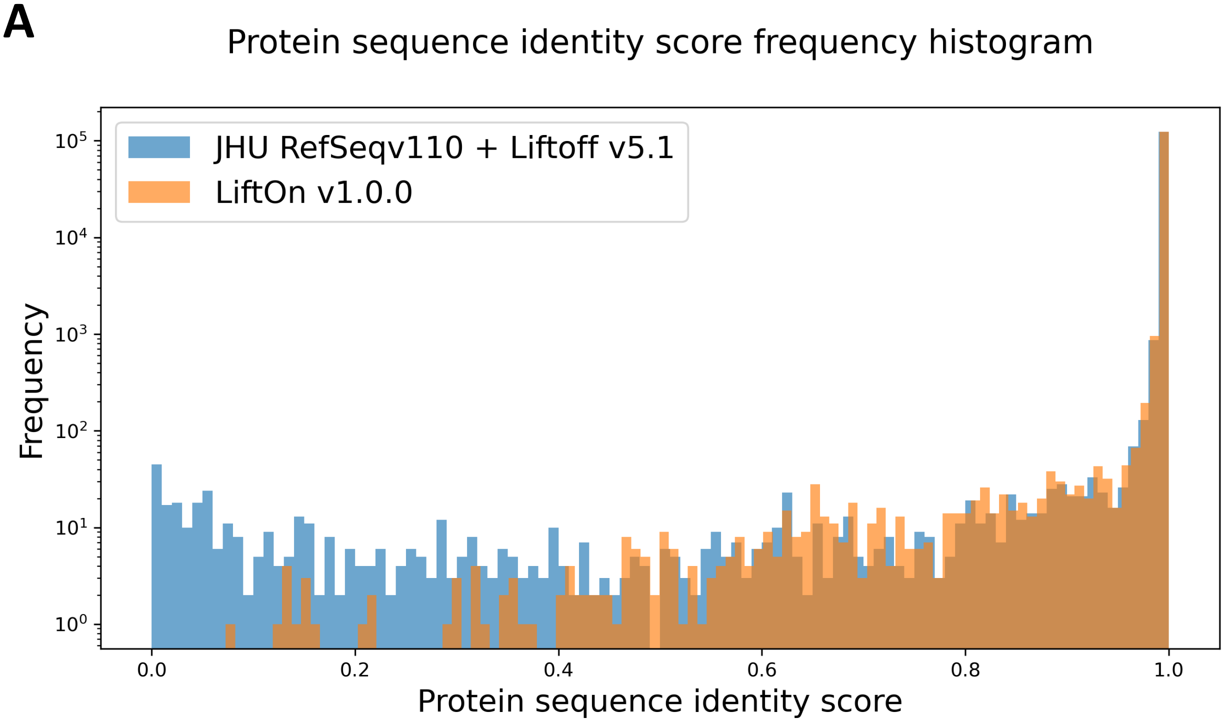


**B**

| | Frameshift | Start_loss | Stop_missing | Stop gain | Inframe insertion | Inframe deletion | Nonsynonymous |
|---|---|---|---|---|---|---|---|
| LiftOn (GRCh38 => CHM13) | 126 | 5 | 106 | 781 | 2 | 21 | 5 |

**Figure S6 (A)** A frequency plot of protein-coding sequence identity scores comparing LiftOn's CHM13 annotation mapped from GRCh38 RefSeq release 220 (yellow) and the published annotation (blue) of CHM13 (JHU RefSeq v110 + Liftoff v5.1, downloaded from https://github.com/marbl/CHM13). **(B)** Count of various mutation types found in transcripts with protein sequence identity score less than 0.95, from the LiftOn mapping of RefSeq from GRCh38 to T2T-CHM13 v2.0.

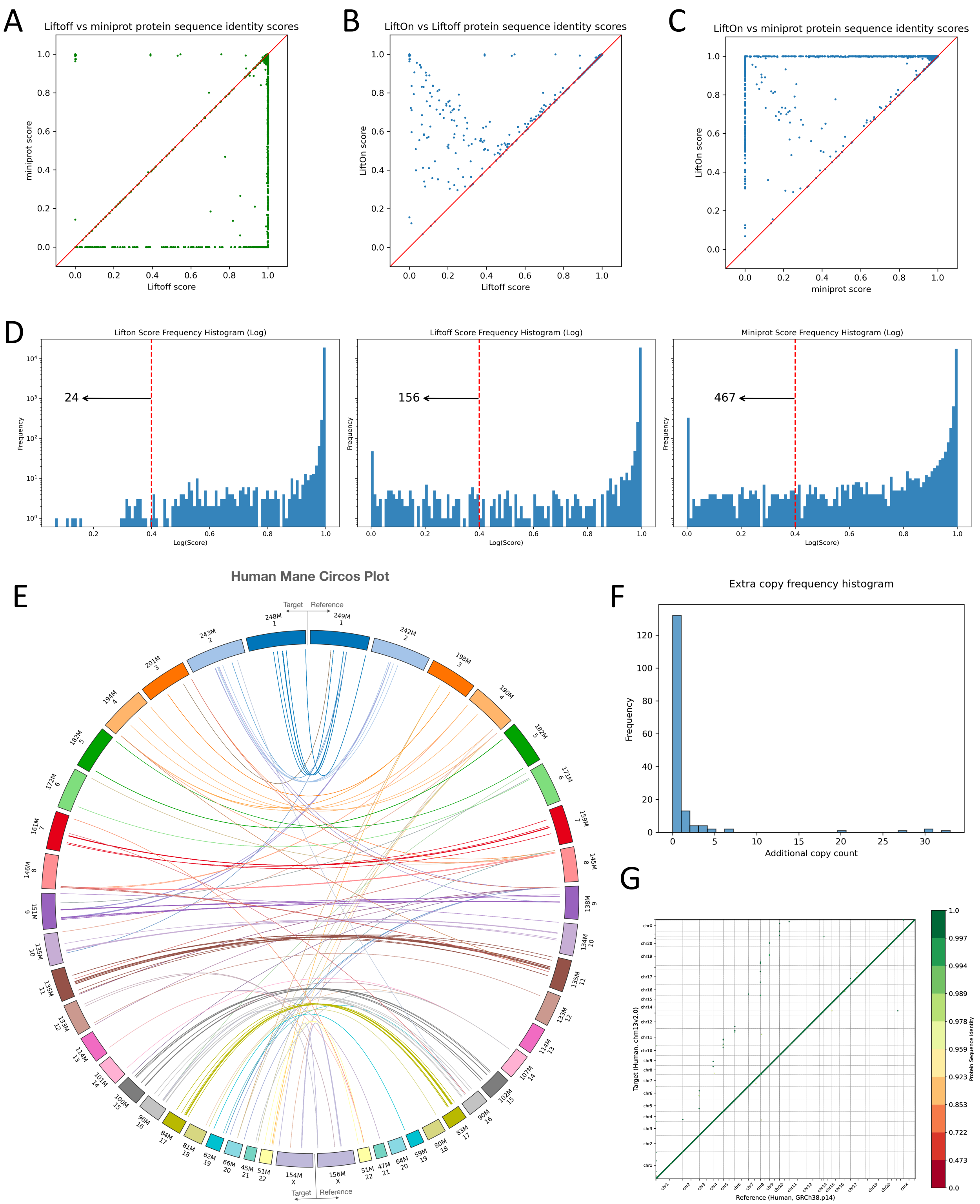


**Figure S7.** Comparative analysis of lifting over RefSeq MANE release v1.2 annotation from GRCh38 to T2T-CHM13. **(A-C)** Scatter plots of protein sequence identity. **(A)** Comparison between miniprot (y-axis) and Liftoff (x-axis), **(B)** comparison between LiftOn (y-axis) and Liftoff (x-axis), and **(C)** comparison between LiftOn (y-axis) and miniprot (x-axis). **(D)** Frequency plots on a logarithmic scale of protein sequence identity for LiftOn (left), Liftoff (middle), and miniprot (right) for the results of GRCh38-to-CHM13 mapping. The score is set to zero for any protein-coding transcript that has not been mapped or mapped at a wrong gene locus. The number of truncated genes with a protein sequence identity lower than 0.4 threshold, as indicated by the red dotted line, is 24 for LiftOn, 156 for Liftoff, and 467 for miniprot. **(E)** Circos plot illustrating the gene loci locations of extra copies found on T2T-CHM13 (left circle) compared to GRCh38 (right circle). Chromosomes of T2T-CHM13 are ordered by the most shared gene copies compared to GRCh38. **(F)** Frequency plot of extra gene copy counts reported by LiftOn. **(G)** Protein-gene order plot, with the x-axis representing the reference genome (GRCh38.p14) and the y-axis representing the target genome (T2T-CHM13v2.0). The protein sequence identities are color-coded on a logarithmic scale, ranging from green to red. Green represents a sequence identity score of 1, while red corresponds to a sequence identity score of 0.

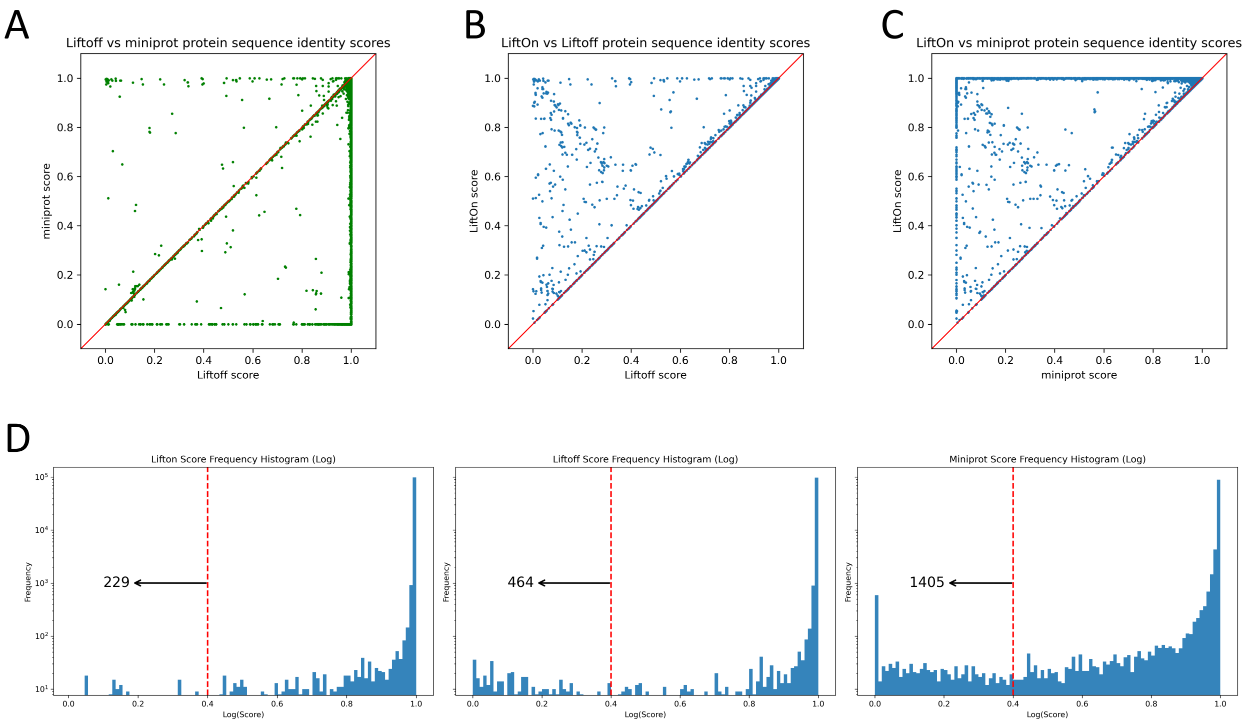


**Figure S8.** Comparative analysis of lifting over CHESS 3 annotation from GRCh38 to T2T-CHM13. **(A-C)** Scatter plots of protein sequence identity. **(A)** Comparison between miniprot (y-axis) and Liftoff (x-axis), **(B)** comparison between LiftOn (y-axis) and Liftoff (x-axis), and **(C)** comparison between LiftOn (y-axis) and miniprot (x-axis). **(D)** Frequency plots on a logarithmic scale of protein sequence identity for LiftOn (left), Liftoff (middle), and miniprot (right) for the results of mouse-to-rat mapping. The score is set to zero for any protein-coding transcript that has not been mapped or mapped at a wrong gene locus. The number of truncated protein-coding transcripts with a protein sequence identity lower than 0.4 threshold, as indicated by the red dotted line, is 229 for LiftOn, 464 for Liftoff, and 1405 for miniprot.

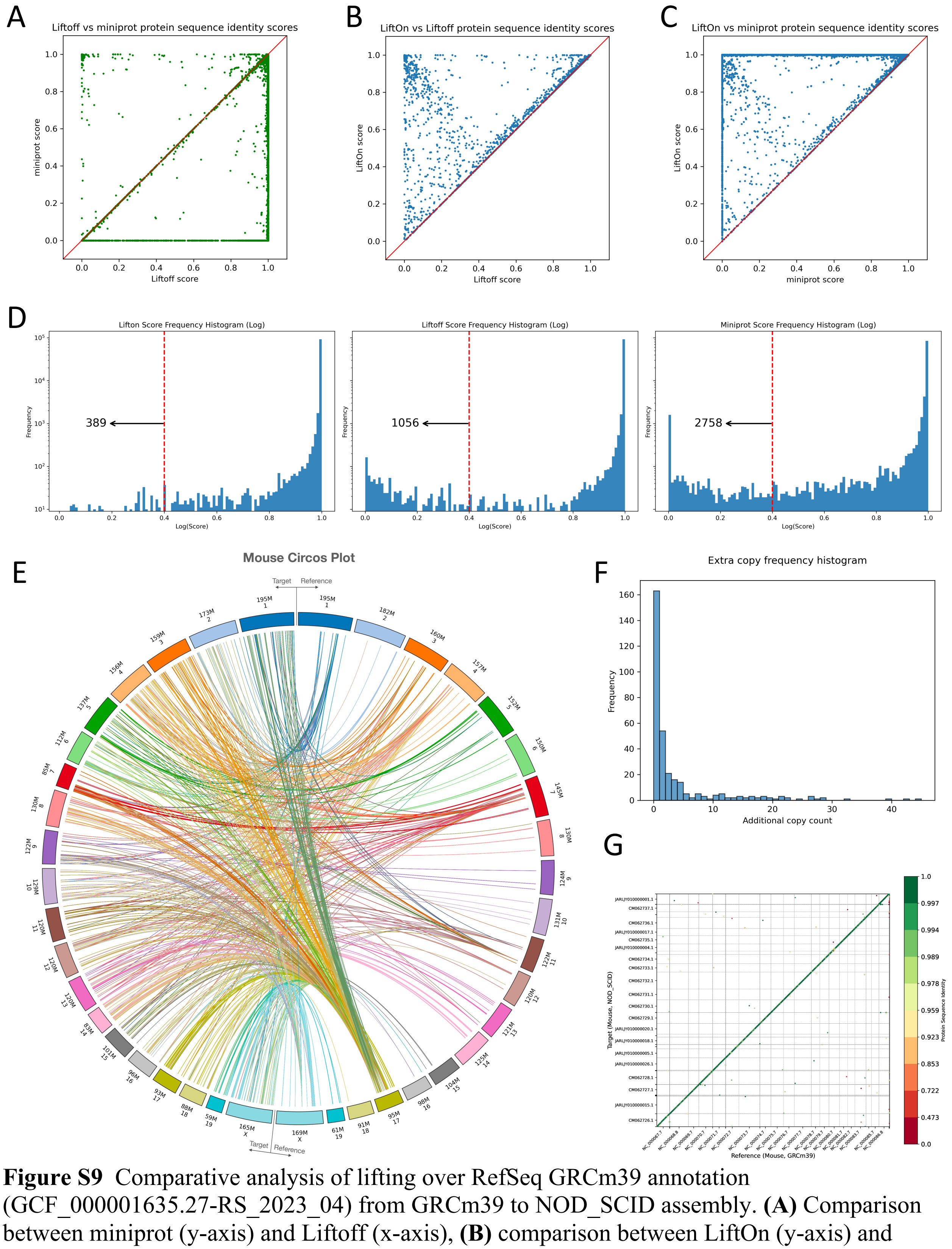


**Figure S9** Comparative analysis of lifting over RefSeq GRCm39 annotation (GCF_000001635.27-RS_2023_04) from GRCm39 to NOD_SCID assembly. **(A)** Comparison between miniprot (y-axis) and Liftoff (x-axis), **(B)** comparison between LiftOn (y-axis) and Liftoff (x-axis), and **(C)** comparison between LiftOn (y-axis) and miniprot (x-axis). **(D)** Frequency plots on a logarithmic scale of protein sequence identity for LiftOn (left), Liftoff (middle), and miniprot (right) for the results of mouse-to-mouse mapping. The score is set to zero for any protein-coding transcript that has not been mapped or mapped at a wrong gene locus. The number of truncated genes with a protein sequence identity lower than 0.4 threshold, as indicated by the red dotted line, is 389 for LiftOn, 1056 for Liftoff, and 2758 for miniprot. **(E)** Circos plot illustrating the gene loci locations of extra copies found on NOD_SCID assembly (left circle) compared to GRCm39 (right circle). Chromosomes of NOD_SCID assembly are ordered by the most shared gene copies compared to GRCm39. **(F)** Frequency plot of extra gene copy counts reported by LiftOn. **(G)** Protein-gene order plot, with the x-axis representing the reference genome (GRCm39) and the y-axis representing the target genome (NOD_SCID assembly). The protein sequence identities are color-coded on a logarithmic scale, ranging from green to red. Green represents a sequence identity score of 1, while red corresponds to a sequence identity score of 0.

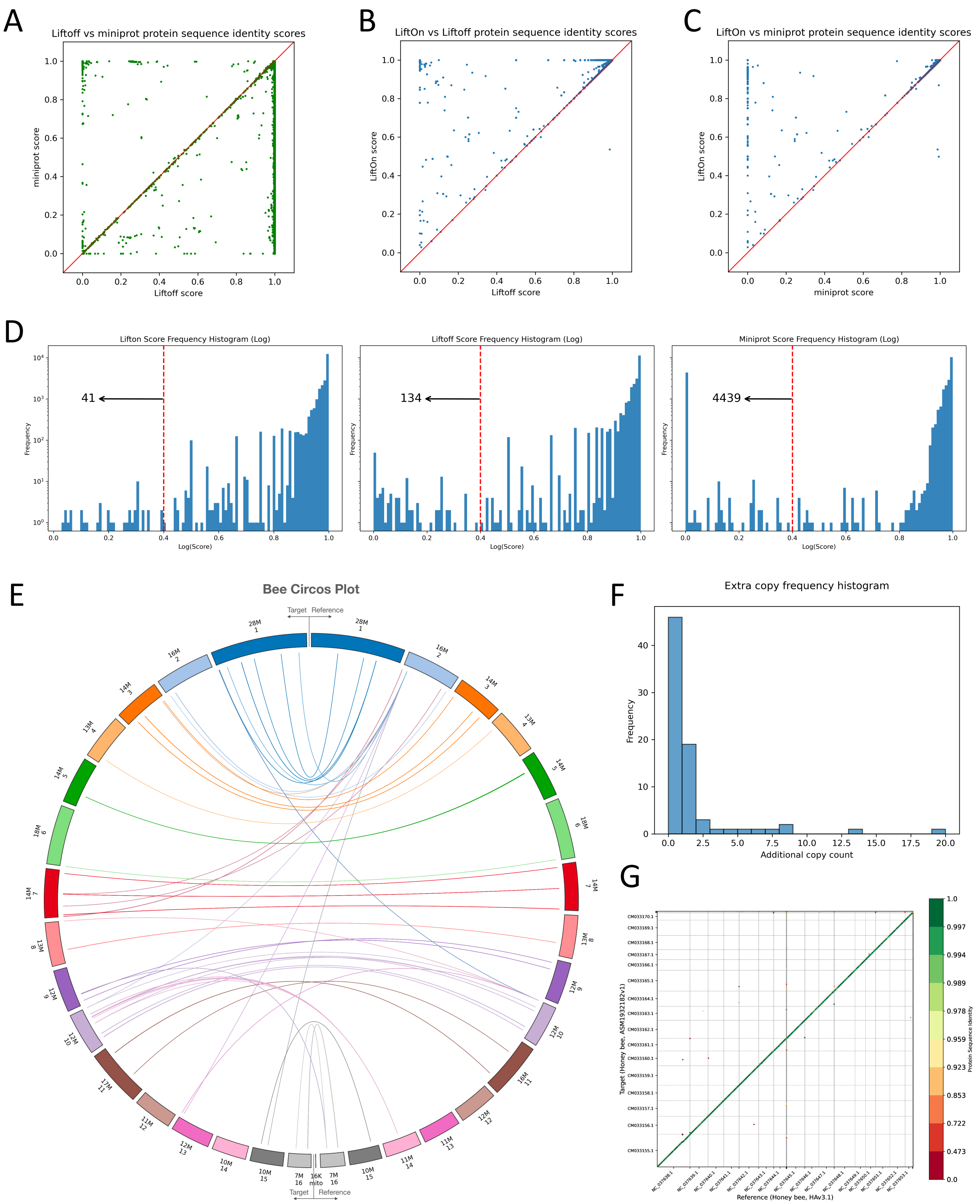


**Figure S10** Comparative analysis of lifting over RefSeq *Apis mellifera* annotation 104 release from Amel_HAv3.1 to ASM1932182v1 assembly. **(A)** Comparison between miniprot (y-axis) and Liftoff (x-axis), **(B)** comparison between LiftOn (y-axis) and Liftoff (x-axis), and **(C)** comparison between LiftOn (y-axis) and miniprot (x-axis). **(D)** Frequency plots on a logarithmic scale of protein sequence identity for LiftOn (left), Liftoff (middle), and miniprot (right) for the results of bee-to-bee mapping. The score is set to zero for any protein-coding transcript that has not been mapped or mapped at a wrong gene locus. The number of truncated genes with a protein sequence identity lower than 0.4 threshold, as indicated by the red dotted line, is 41 for LiftOn, 134 for Liftoff, and 4439 for miniprot. **(E)** Circos plot illustrating the gene loci locations of extra copies found on ASM1932182v1 assembly (left circle) compared to Amel_HAv3.1 (right circle). Chromosomes of ASM1932182v1 assembly are ordered by the most shared gene copies compared to Amel_HAv3.1. **(F)** Frequency plot of extra gene copy counts reported by LiftOn. **(G)** Protein-gene order plot, with the x-axis representing the reference genome (Amel_HAv3.1) and the y-axis representing the target genome (ASM1932182v1 assembly). The protein sequence identities are color-coded on a logarithmic scale, ranging from green to red. Green represents a sequence identity score of 1, while red corresponds to a sequence identity score of 0.

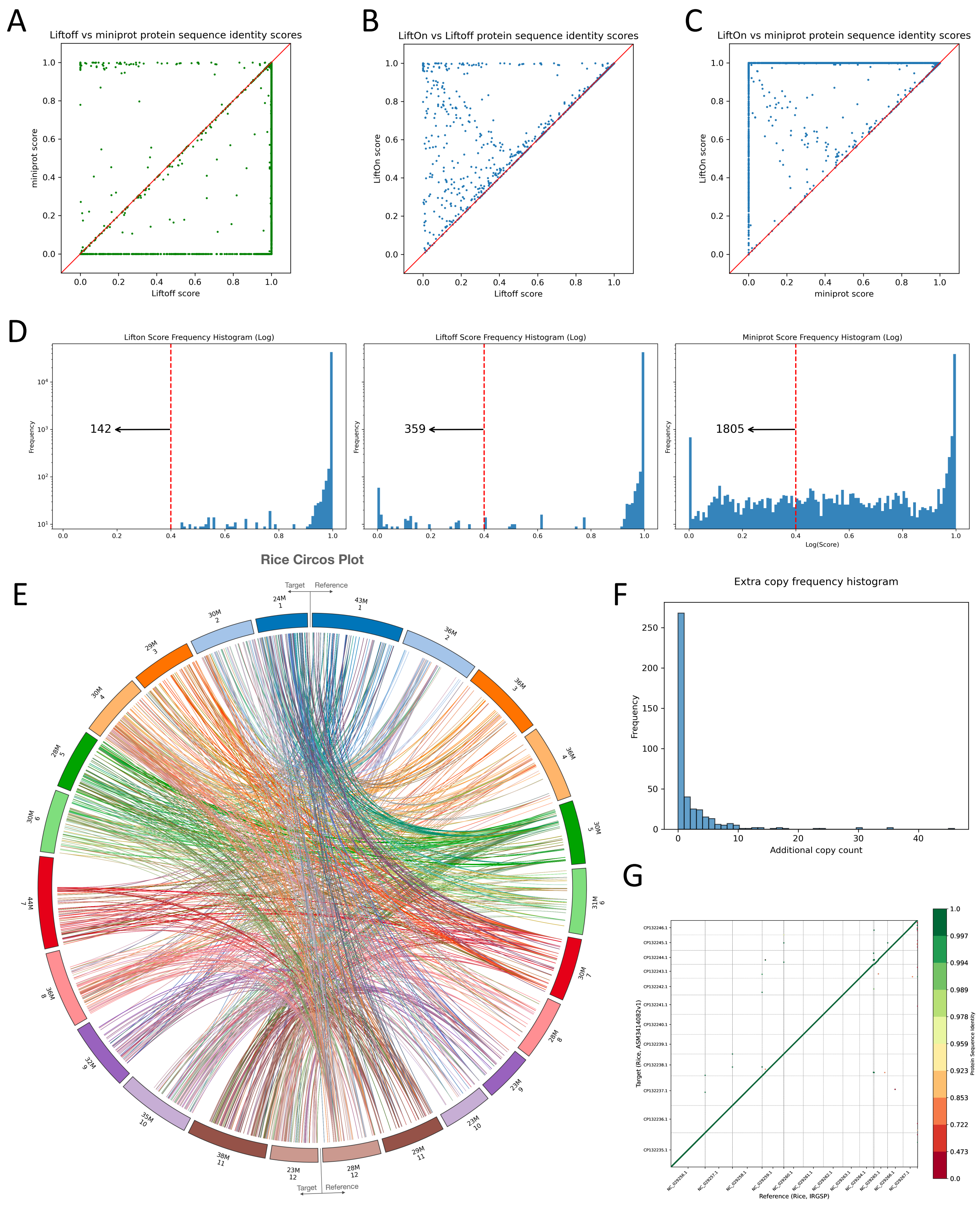


**Figure S11** Comparative analysis of lifting over RefSeq annotation 102 release from IRGSP-1.0 to ASM3414082v1 assembly. **(A)** Comparison between miniprot (y-axis) and Liftoff (x-axis), **(B)** comparison between LiftOn (y-axis) and Liftoff (x-axis), and **(C)** comparison between LiftOn (y-axis) and miniprot (x-axis). **(D)** Frequency plots on a logarithmic scale of protein sequence identity for LiftOn (left), Liftoff (middle), and miniprot (right) for the results of rice-to-rice mapping. The score is set to zero for any protein-coding transcript that has not been mapped or mapped at a wrong gene locus. The number of truncated genes with a protein sequence identity lower than 0.4 threshold, as indicated by the red dotted line, is 142 for LiftOn, 359 for Liftoff, and 1805 for miniprot. **(E)** Circos plot illustrating the gene loci locations of extra copies found on ASM3414082v1 assembly (left circle) compared to IRGSP-1.0 (right circle). Chromosomes of ASM3414082v1 assembly are ordered by the most shared gene copies compared to IRGSP-1.0. **(F)** Frequency plot of extra gene copy counts reported by LiftOn. **(G)** Protein-gene order plot, with the x-axis representing the reference genome (IRGSP-1.0) and the y-axis representing the target genome (ASM3414082v1 assembly). The protein sequence identities are color-coded on a logarithmic scale, ranging from green to red. Green represents a sequence identity score of 1, while red corresponds to a sequence identity score of 0.

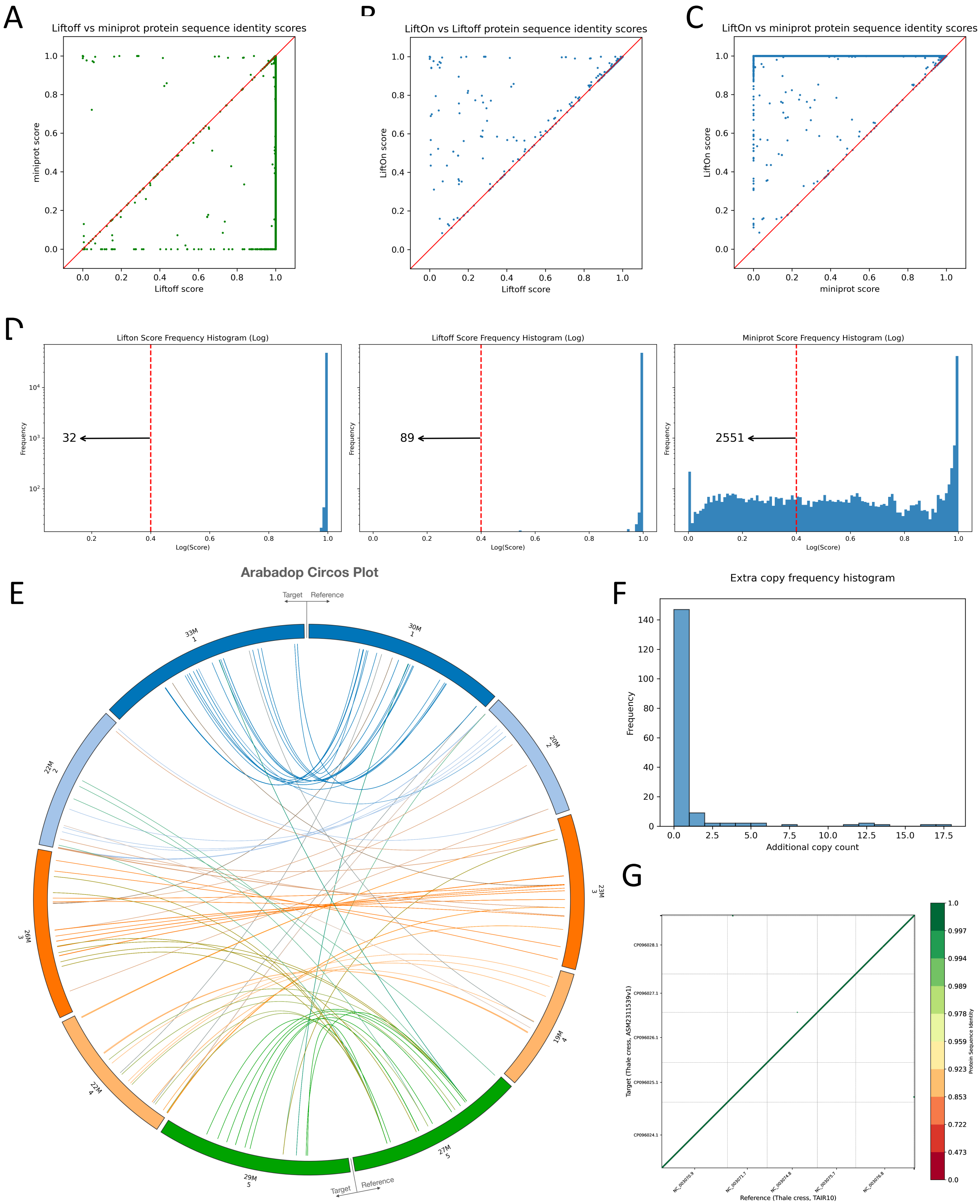


**Figure S12** Comparative analysis of lifting over RefSeq annotations from TAIR10.1 to ASM2311539v1. **(A)** Comparison between miniprot (y-axis) and Liftoff (x-axis), **(B)** comparison between LiftOn (y-axis) and Liftoff (x-axis), and **(C)** comparison between LiftOn (y-axis) and miniprot (x-axis). **(D)** Frequency plots on a logarithmic scale of protein sequence identity for LiftOn (left), Liftoff (middle), and miniprot (right) for the results of arabidopsis-to-arabidopsis mapping. The score is set to zero for any protein-coding transcript that has not been mapped or mapped at a wrong gene locus. The number of truncated genes with a protein sequence identity lower than 0.4 threshold, as indicated by the red dotted line, is 32 for LiftOn, 89 for Liftoff, and 2551 for miniprot. **(E)** Circos plot illustrating the gene loci locations of extra copies found on ASM2311539v1 assembly (left circle) compared to TAIR10.1 (right circle). Chromosomes of ASM2311539v1 assembly are ordered by the most shared gene copies compared to TAIR10.1. **(F)** Frequency plot of extra gene copy counts reported by LiftOn. **(G)** Protein-gene order plot, with the x-axis representing the reference genome (TAIR10.1) and the y-axis representing the target genome (ASM2311539v1 assembly). The protein sequence identities are color-coded on a logarithmic scale, ranging from green to red. Green represents a sequence identity score of 1, while red corresponds to a sequence identity score of 0.

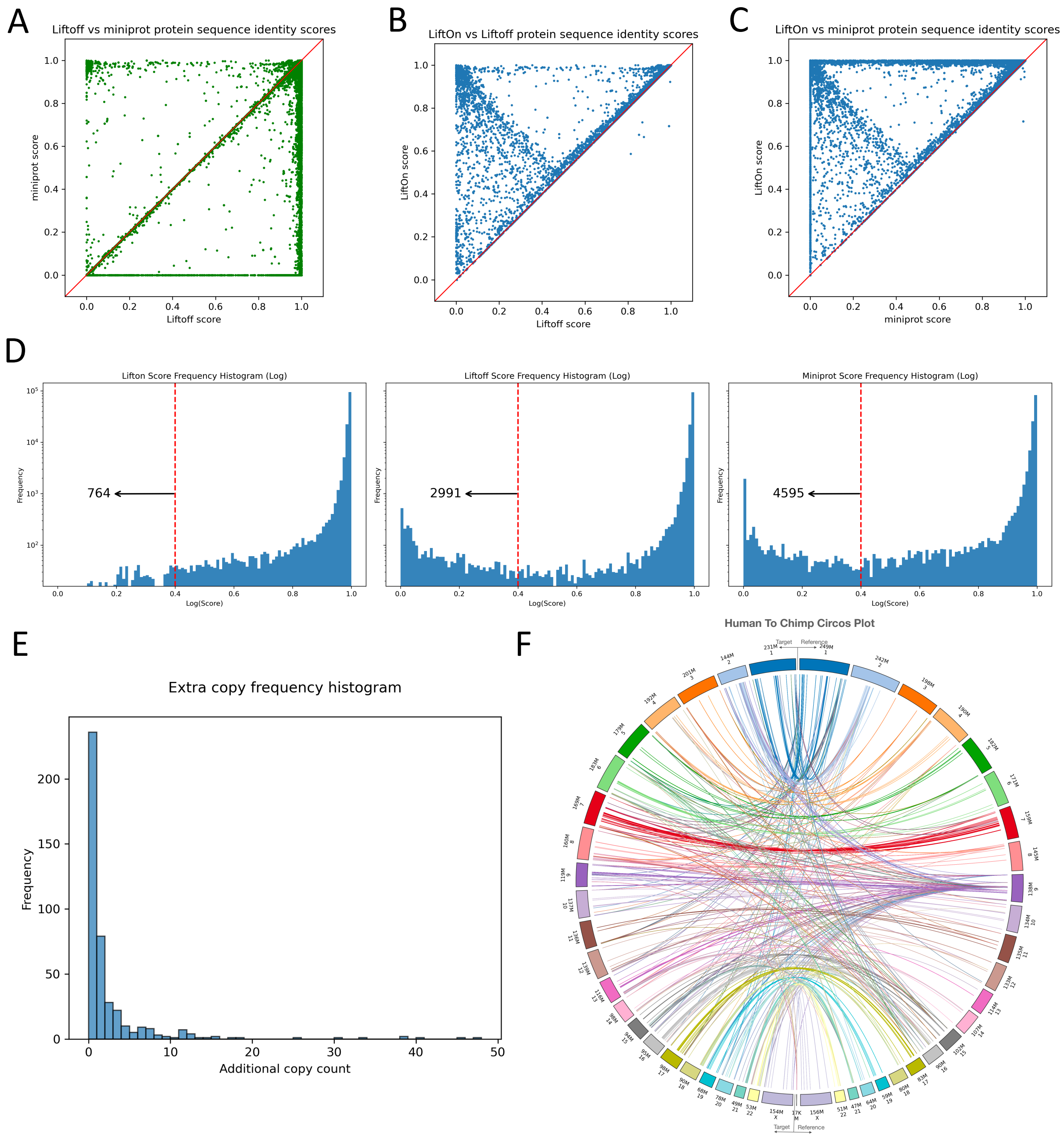


**Figure S13** Comparative analysis of lifting over RefSeq release GCF_000001405.40-RS_2023_10 of GRCh38.p14 from GRCh38 to NHGRI_mPanTro3-v1.1. **(A)** Comparison between miniprot (y-axis) and Liftoff (x-axis), **(B)** comparison between LiftOn (y-axis) and Liftoff (x-axis), and **(C)** comparison between LiftOn (y-axis) and miniprot (x-axis). **(D)** Frequency plots on a logarithmic scale of protein sequence identity for LiftOn (left), Liftoff (middle), and miniprot (right) for the results of human-to-chimpanzee mapping. The score is set to zero for any protein-coding transcript that has not been mapped or mapped at a wrong gene locus. The number of truncated genes with a protein sequence identity lower than 0.4 threshold, as indicated by the red dotted line, is 764 for LiftOn, 2991 for Liftoff, and 4595 for miniprot. **(E)**

Frequency plot of extra gene copy counts reported by LiftOn. **(F)** Circos plot illustrating the gene loci locations of extra copies found on NHGRI_mPanTro3-v1.1 assembly (left circle) compared to GRCh38 (right circle). Chromosomes of NHGRI_mPanTro3-v1.1 assembly are ordered by the most shared gene copies compared to GRCh38.

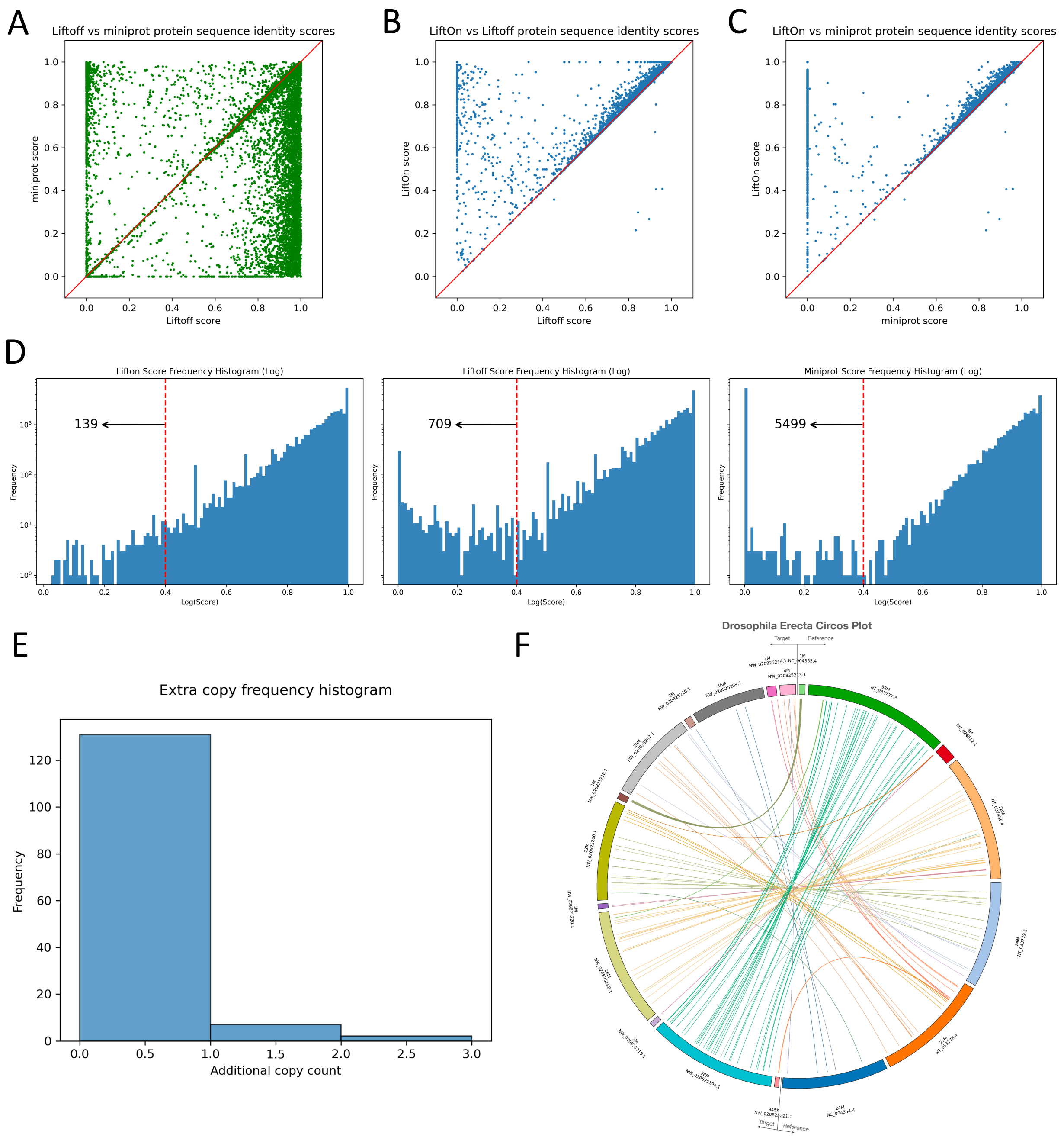


**Figure S14** Comparative analysis of lifting over RefSeq FlyBase Release 6.54 annotations from Release 6 + ISO1 MT (*Drosophila melanogaster*) to DereRS2 (*Drosophila erecta*). **(B)** comparison between LiftOn (y-axis) and Liftoff (x-axis), and **(C)** comparison between LiftOn (y-axis) and miniprot (x-axis). **(D)** Frequency plots on a logarithmic scale of protein sequence identity for LiftOn (left), Liftoff (middle), and miniprot (right) for the results of mouse-to-rat mapping. The score is set to zero for any protein-coding transcript that has not been mapped or mapped at a wrong gene locus. The number of truncated genes with a protein sequence identity lower than 0.4 threshold, as indicated by the red dotted line, is 139 for LiftOn, 709 for Liftoff, and 5499 for miniprot. **(E)** Frequency plot of extra gene copy counts reported by LiftOn. **(F)** Circos plot illustrating the gene loci locations of extra copies found on DereRS2 assembly (left circle)

compared to Release 6 + ISO1 MT (right circle). Chromosomes of DereRS2 assembly are ordered by the most shared gene copies compared to Release 6 + ISO1 MT.

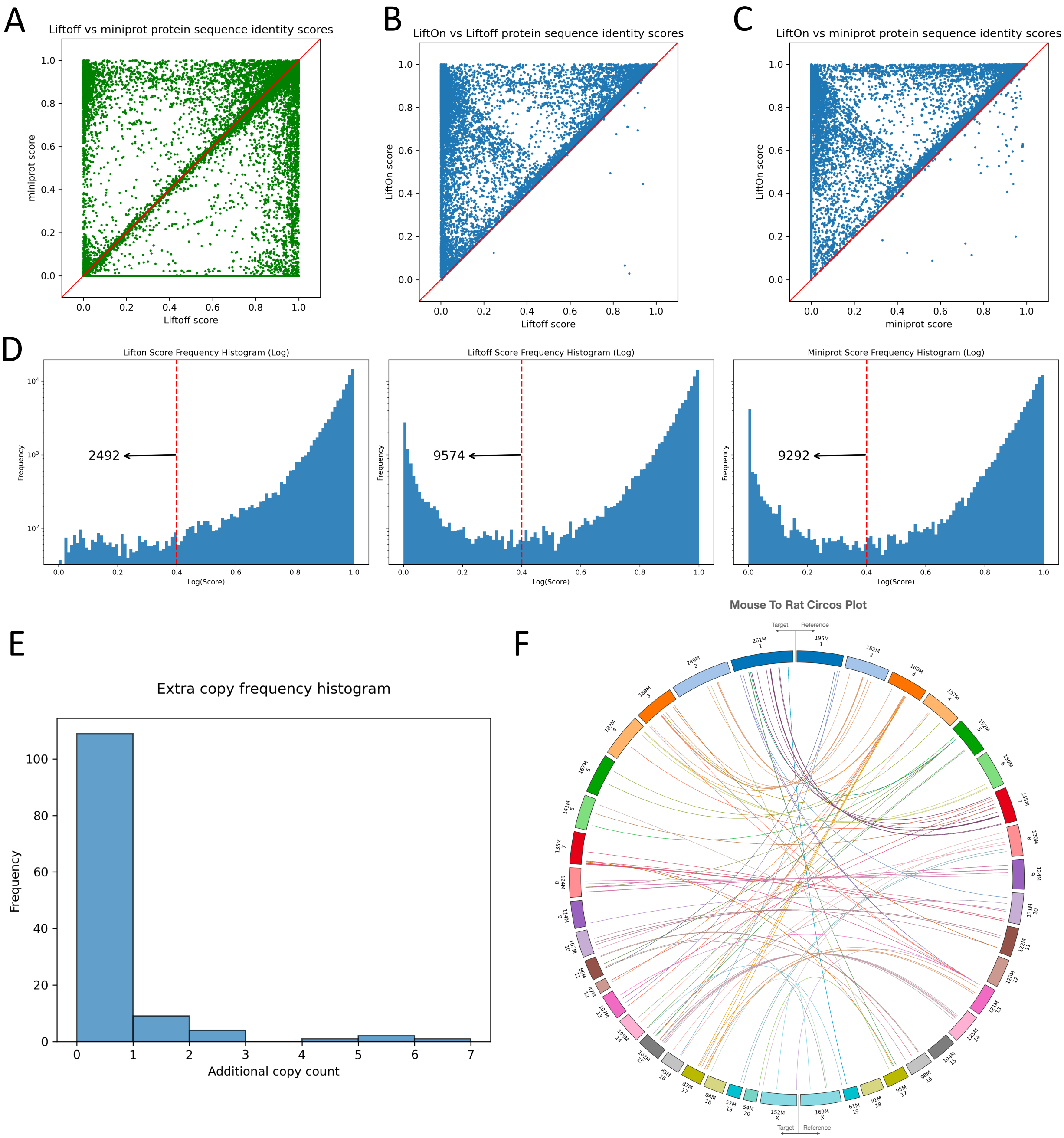


**Figure S15** Comparative analysis of lifting over GRCm39 annotation (GCF_000001635.27-RS_2023_04) from GRCm39 (*Mus musculus*) to mRatBN7.2 (*Rattus norvegicus*). **(A)** Comparison between miniprot (y-axis) and Liftoff (x-axis), **(B)** comparison between LiftOn (y-axis) and Liftoff (x-axis), and **(C)** comparison between LiftOn (y-axis) and miniprot (x-axis). **(D)** Frequency plots on a logarithmic scale of protein sequence identity for LiftOn (left), Liftoff (middle), and miniprot (right) for the results of mouse-to-rat mapping. The score is set to zero for any protein-coding transcript that has not been mapped or mapped at a wrong gene locus. The number of truncated genes with a protein sequence identity lower than 0.4 threshold, as indicated by the red dotted line, is 2492 for LiftOn, 9574 for Liftoff, and 9292 for miniprot. **(E)** Frequency plot of extra gene copy counts reported by LiftOn. **(F)** Circos plot illustrating the gene loci locations of extra copies found on mRatBN7.2 assembly (left circle) compared to GRCm39 (right circle). Chromosomes of mRatBN7.2 assembly are ordered by the most shared gene copies compared to GRCm39.

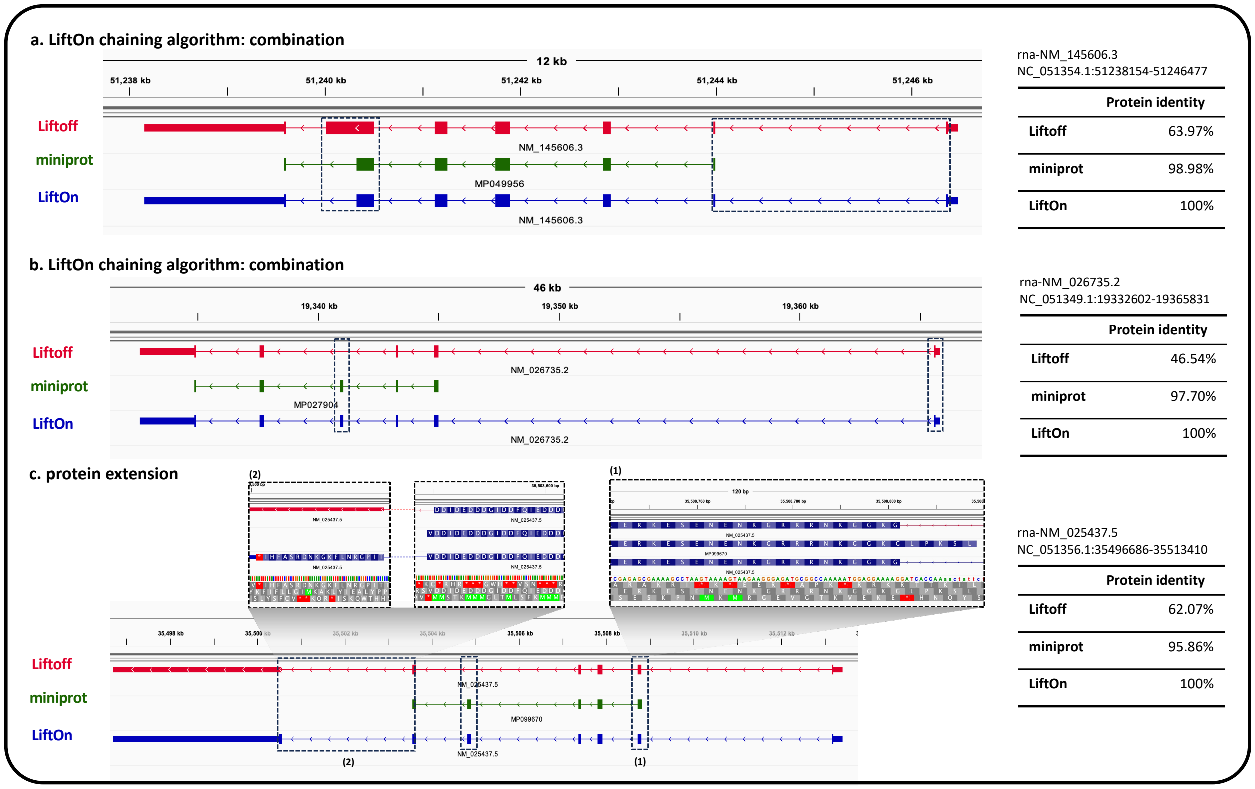


**Figure S16** Selected examples from IGV demonstrate LiftOn's ability to generate novel protein-coding transcripts, surpassing Liftoff and miniprot readouts. **(A)** demonstrates LiftOn correctly locating the starting region of a short CDS which miniprot misses (producing subsequent error in the next CDS) but Liftoff catches, and a stop codon in a CDS which Liftoff misses but miniprot catches, yielding a consensus with 100% protein identity score. Similarly, **(B)** demonstrates LiftOn catching the short starting CDS which miniprot misses but Liftoff catches, and a missing CDS which Liftoff misses but miniport catches. Lastly, **(C)** again showcases LiftOn producing the best consensus between Liftoff and miniprot, while also highlighting a new feature in (2), where the algorithm extends the protein with an open reading frame search for a novel terminal CDS, yielding the complete protein sequence which both Liftoff and miniprot miss.

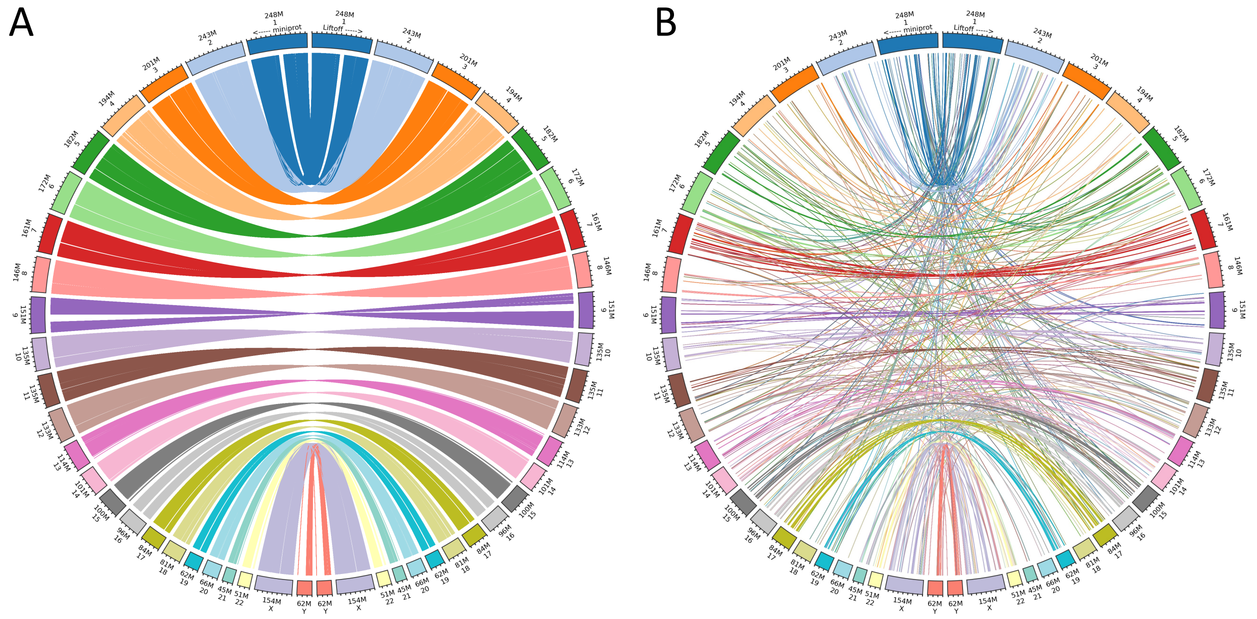


**Figure S17** Circos plots illustrating the comparison of protein-coding transcript coordinates between Liftoff and miniprot annotations based on the mapping of RefSeq annotation v220 from GRCh38 to T2T-CHM13, as described in Results 2.2. (A) Circos plot displays protein-coding transcript coordinates where miniprot aligns proteins to the same gene loci as Liftoff. (B) Circos plot depicts protein-coding transcript coordinates where miniprot aligns proteins to different gene loci than Liftoff. In both circos plots, the left circle represents miniprot coordinates on T2T-CHM13, while the right circle shows Liftoff coordinates on T2T-CHM13.

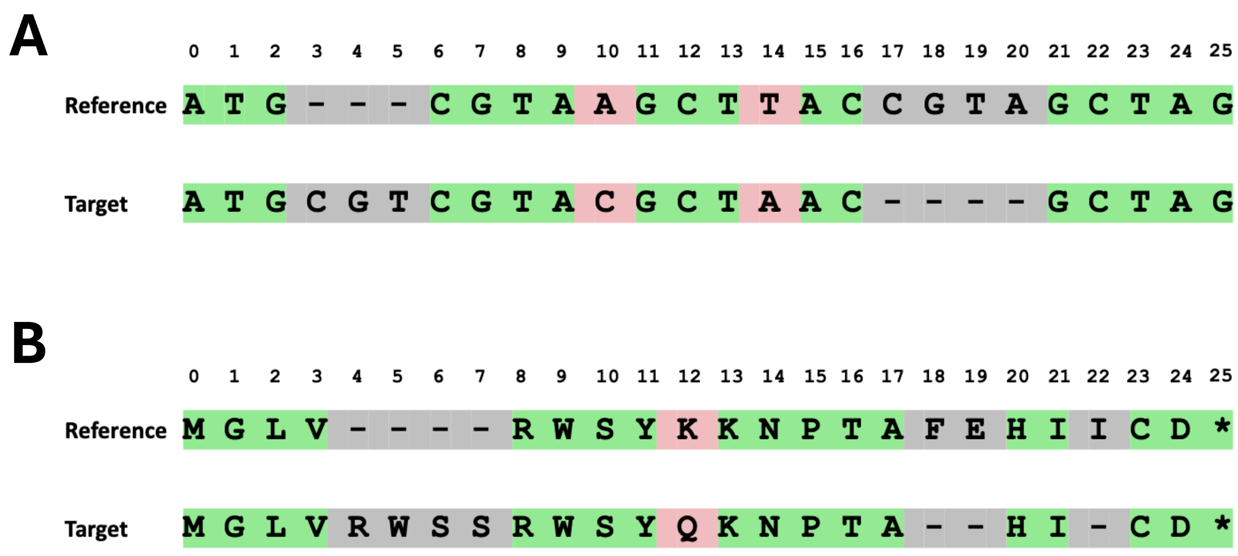


**Figure S18** Examples of calculating **(A)** DNA and **(B)** protein sequence identity scores. **(A)** Alignment of reference and target DNA sequences, with 17 out of 26 alignment columns containing matched nucleotides, results in a DNA sequence identity score of 65.4% ($\frac{\#Matched\_nucleotide}{\#\ alignment\_column}$). **(B)** Alignment of reference and target protein sequences, with 18 out of 26 alignment columns containing matched amino acids and 4 gaps in the reference, results in a protein sequence identity score of 81.8% ($\frac{\#Matched\_AA}{\#\ alignment\_column\ - \#gaps\_in\_reference\_protein}$).
