## Supplementary Tables for "Combining DNA and protein alignments to improve genome annotation with LiftOn"

| Gene biotype | Number of genes in GRCh38 | Number of genes mapped onto T2T_CHM13 by LiftOn |
|---:|---:|---:|
| protein_coding | 19927 | 20144 |
| lncRNA | 18008 | 18723 |
| Pseudogene | 16896 | 17855 |
| miRNA | 1915 | 2383 |
| snoRNA | 1194 | 1187 |
| tRNA | 453 | 548 |
| V_segment | 239 | 244 |
| snRNA | 153 | 192 |
| J_segment | 98 | 80 |
| ncRNA | 51 | 49 |
| misc_RNA | 42 | 44 |
| C_region | 21 | 23 |
| antisense_RNA | 19 | 19 |
| other | 13 | 13 |
| Y_RNA | 4 | 7 |
| vault_RNA | 4 | 4 |
| scRNA | 4 | 4 |
| telomerase_RNA | 1 | 1 |
| RNase_P_RNA | 1 | 1 |
| RNase_MRP_RNA | 1 | 1 |
| **Total** | **59044** | **61522** |

**Table S1** The number of genes for each biotype annotated on the main chromosomes and unplaced contigs in GRCh38 and T2T-CHM13. The T2T-CHM13 annotations were generated using LiftOn by mapping the RefSeq annotations (Release 220) from GRCh38 patch 14 to T2T-CHM13 v2.0.

| Gene Name | Number of extra copies |
|---|---|
| gene-LOC124905331 | 50 |
| gene-TAF11L5 | 34 |
| gene-USP17L11 | 25 |
| gene-TSPY10 | 19 |
| gene-FAM90A13 | 14 |
| gene-FAM90A12 | 9 |
| gene-LOC101929601 | 7 |
| gene-FAM90A24, gene-TSPY4 | 6 |
| gene-TSPY3 | 5 |
| gene-AMY1C, gene-LOC124900996, gene-LOC124901652, gene-LOC124903857 | 4 |
| gene-LOC107987371, gene-LOC124901646, gene-LOC124903544, gene-TSPY9, gene-USP17L17 | 3 |
| gene-CLEC18B, gene-GOLGA6B, gene-LOC107987020, gene-LOC107987067, gene-LOC107987372, gene-LOC112268317, gene-LOC124900992, gene-LOC124901580, gene-LOC124901639, gene-LOC124901648, gene-LOC124901651, gene-LOC124901712, gene-LOC124904581, gene-LOC124905938, gene-LOC128966684, gene-NPIPB15, gene-PDPR, gene-PGA4, gene-TSPY8 | 2 |
| gene-BOLA2, gene-CCL3L3, gene-CCL4L2, gene-DEFB103B, gene-DEFB104A, gene-DEFB105B, gene-DEFB106A, gene-DEFB107A, gene-DEFB4A, gene-DUSP22, gene-EIF3C, gene-FAM90A9, gene-FCGR3B, gene-FOXD4L4, gene-FRG2C, gene-GOLGA6L1, gene-GOLGA8R, gene-GPRIN2, gene-KCNJ18, gene-LGALS9C, gene-LIMS3, gene-LOC101927345, gene-LOC101929627, gene-LOC107985915, gene-LOC112268458, gene-LOC124901638, gene-LOC124901798, gene-LOC124903442, gene-LOC124903761, gene-LOC124904095, gene-LOC124905300, gene-LOC124905320, gene-LOC124907854, gene-LOC128966594, gene-MRGPRX1, gene-PRR23D2, gene-RGPD5, gene-SLX1A, gene-SPAG11B, gene-SPDYE13, gene-SPDYE9, gene-SULT1A3, gene-TPTE, gene-TSPY1, gene-USP17L7, gene-XAGE1A, gene-ZNG1C | 1 |

**Table S2.** Distribution of extra copies of protein-coding genes identified by the Liftoff module in LiftOn of mapping RefSeq GCF_000001405.40-RS_2023_10 release annotations from GRCh38.p14 to T2T-CHM13 v2.0.

| Gene Name | Number of extra copies |
|---|---|
| gene-LOC112268317 | 27 |
| gene-LOC124903544, gene-LOC124902738, gene-IGHJ6, gene-KCNE1 | 1 |

**Table S3.** Distribution of extra copies of protein-coding genes identified by the miniprot module in LiftOn of mapping RefSeq GCF_000001405.40-RS_2023_10 release annotations from GRCh38.p14 to T2T-CHM13 v2.0.

| | Total gene count | Protein-coding gene count | | | Non-coding gene count | | |
|---|---|---|---|---|---|---|---|
| *Reference* | 19,158 | 19,158 | | | 0 | | |
| *Target (LiftOn)* | 19,535 | *Single copy* | *Extra copy* | *Extra copy count* | *Single copy* | *Extra copy* | *Extra copy count* |
| | | 18,968 | 162 | 405 | 0 | 0 | 0 |
| | | 19,535 | | | 0 | | |

**Table S4.** Statistics for LiftOn at the gene level, as a result of mapping RefSeq MANE release v1.2 annotation from the GRCh38.p14 human genome to T2T-CHM13 v2.0 (https://github.com/marbl/CHM13).

| | | Total feature count | Protein-coding feature count | | | Non-coding feature count | | |
|---|---|---|---|---|---|---|---|---|
| Transcript | Reference | 168,451 | 105,328 | | | 69,250 | | |
| | Target (LiftOn) | 156,173 | *Single copy* | *Extra copy* | *Extra copy count* | *Single copy* | *Extra copy* | *Extra copy count* |
| | | | 99,461 | 45 | 138 | 55,217 | 139 | 1,357 |
| | | | 99,599 | | | 56,574 | | |

**Table S5.** The summary LiftOn statistics for lift-over results at transcript-levels, depicting the mapping from Human GRCh38 CHESS 3 v .3.0.1 (https://ccb.jhu.edu/chess/) to T2T-CHM13 v2.0 (https://github.com/marbl/CHM13)

| | | Total feature count | Protein-coding feature count | | | Non-coding feature count | | |
|---|---|---|---|---|---|---|---|---|
| Gene | Reference | 35,551 | 22,192 | | | 13,359 | | |
| Gene | Target (LiftOn) | 36,525 | *Single copy* | *Extra copy* | *Extra copy count* | *Single copy* | *Extra copy* | *Extra copy count* |
| | | | 21,507 | 198 | 840 | 13,195 | 131 | 654 |
| | | | 22,545 | | | 13,980 | | |
| Transcript | Reference | 119,745 | 96,192 | | | 23,553 | | |
| Transcript | Target (LiftOn) | 120,692 | *Single copy* | *Extra copy* | *Extra copy count* | *Single copy* | *Extra copy* | *Extra copy count* |
| | | | 95,169 | 280 | 986 | 23,284 | 173 | 800 |
| | | | 96,435 | | | 24,257 | | |

**Table S6** Statistics for LiftOn at both the gene and transcript levels, as a result of mapping the RefSeq GRCm39 annotation (GCF_000001635.27-RS_2023_04) from GRCm39 (C57BL/6J strain) to NOD_SCID assembly (NOD_SCID strain).

<table>
<tr><th></th><th></th><th>Total feature count</th><th colspan="3">Protein-coding feature count</th><th colspan="3">Non-coding feature count</th></tr>
<tr><td rowspan="4">Gene</td><td>Reference</td><td>11,793</td><td colspan="3">9,935</td><td colspan="3">1,858</td></tr>
<tr><td rowspan="3">Target (LiftOn)</td><td rowspan="3">11,916</td><td>Single copy</td><td>Extra copy</td><td>Extra copy count</td><td>Single copy</td><td>Extra copy</td><td>Extra copy count</td></tr>
<tr><td>9,841</td><td>54</td><td>95</td><td>1,822</td><td>23</td><td>81</td></tr>
<tr><td colspan="3">9,990</td><td colspan="3">1,926</td></tr>
<tr><td rowspan="4">Transcript</td><td>Reference</td><td>26,617</td><td colspan="3">23,471</td><td colspan="3">3,146</td></tr>
<tr><td rowspan="3">Target (LiftOn)</td><td rowspan="3">26,797</td><td>Single copy</td><td>Extra copy</td><td>Extra copy count</td><td>Single copy</td><td>Extra copy</td><td>Extra copy count</td></tr>
<tr><td>23,287</td><td>83</td><td>144</td><td>3,078</td><td>45</td><td>160</td></tr>
<tr><td colspan="3">23,514</td><td colspan="3">3,283</td></tr>
</table>

**Table S7** Statistics for LiftOn at both the gene and transcript levels, as a result of mapping the RefSeq *Apis mellifera* annotation release 104 from Amel_HAv3.1 (GCF_003254395.2) to ASM1932182v1 assembly (GCA_019321825.1).

| | | Total feature count | Protein-coding feature count | | | Non-coding feature count | | |
|---|---|---|---|---|---|---|---|---|
| Gene | Reference | 32,008 | 28,738 | | | 3,270 | | |
| | Target (LiftOn) | 33,450 | *Single copy* | *Extra copy* | *Extra copy count* | *Single copy* | *Extra copy* | *Extra copy count* |
| | | | 28,385 | 322 | 1039 | 3,157 | 108 | 439 |
| | | | 29,746 | | | 3,704 | | |
| Transcript | Reference | 48,843 | 42,566 | | | 6,277 | | |
| | Target (LiftOn) | 50,352 | *Single copy* | *Extra copy* | *Extra copy count* | *Single copy* | *Extra copy* | *Extra copy count* |
| | | | 42,181 | 346 | 1047 | 6,146 | 125 | 507 |
| | | | 43,574 | | | 6,778 | | |

**Table S8** Statistics for LiftOn at both the gene and transcript levels, as a result of mapping the NCBI RefSeq *Oryza sativa* Japonica Group annotation release 102 from IRGSP-1.0 (GCF_001433935.1) to ASM3414082v1 assembly (GCA_034140825.1).

| | | Total feature count | Protein-coding feature count | | | Non-coding feature count | | |
| --- | --- | --- | --- | --- | --- | --- | --- | --- |
| Gene | Reference | 31,481 | 27,562 | | | 3,919 | | |
| | Target (LiftOn) | | *Single copy* | *Extra copy* | *Extra copy count* | *Single copy* | *Extra copy* | *Extra copy count* |
| | | 31,525 | 27,390 | 134 | 224 | 3,665 | 38 | 74 |
| | | | 27,748 | | | 3,777 | | |
| Transcript | Reference | 52,635 | 48,256 | | | 4,379 | | |
| | Target (LiftOn) | | *Single copy* | *Extra copy* | *Extra copy count* | *Single copy* | *Extra copy* | *Extra copy count* |
| | | 52,680 | 48,060 | 146 | 237 | 4,125 | 38 | 74 |
| | | | 48,443 | | | 4,237 | | |

**Table S9** Statistics for LiftOn at both the gene and transcript levels, as a result of mapping the RefSeq TAIR10.1 annotation from TAIR10.1 (GCF_000001735.4) to Col-CEN (ASM2311539v1) assembly (GCA_023115395.1).
