## Supplementary Algorithms for "Combining DNA and protein alignments to improve genome annotation with LiftOn"

**Algorithm S1:** The protein-maximization algorithm pairing Liftoff and miniprot gene loci

```
# Inputs:
#   - lifton_gene: An instance of the LiftOn class, which stores gene information.
#   - locus: A Feature instance from the gffutils package representing a genomic locus.
#   - ref_db, liftoff_db, miniprot_db: FeatureDB instances from the gffutils package for reference,
#     liftoff, and miniprot annotations, respectively.
#   - m_id_trans_dict: Dictionary mapping reference transcript IDs to lists of miniprot transcript IDs.
#   - tree_dict: Dictionary mapping reference chromosome IDs to IntervalTree instances.
#   - tgt_fai: Fasta instance from the pyfaidx package, storing target genome sequences.
#   - ref_proteins, ref_trans: Fasta instances from the pyfaidx package, storing reference protein
#     and transcript sequences, respectively.
#   - features_dict: Dictionary mapping reference feature IDs to Lifton_feature instances.
#   - ENTRY_FEATURE: Boolean flag indicating if the current feature is the root feature for lifting over.
# Returns:
#   - lifton_gene: Updated or new lifton_gene instance with processed data.

# Process liftoff annotation and create lifton_gene instance
function protein_maximization(lifton_gene, locus, ref_db, liftoff_db, m_id_trans_dict, miniprot_db,
                              tree_dict, tgt_fai, ref_proteins, ref_trans, features_dict, ENTRY_FEATUR=False):
    exon children = liftoff_db.children(locus, featuretype='exon', level=1, order_by='start')
    if lifton_gene is None and ENTRY_FEATURE:  # LiftOn gene initialization
        lifton_gene, ref_gene_id, ref_trans_id =initialize_lifton_gene(locus, ref_db, tree_dict, features_dict)
        if lifton_gene.ref_gene_id is None: return None

    if len(exon_children) == 0:  # processing features without exons
        parent_feature = lifton_gene if ENTRY_FEATURE else lifton_gene.add_feature(deepcopy(locus))
        features = liftoff_db.children(locus, level=1)
        for feature in features:
            lifton_gene = protein_maximization(parent_feature, feature, ref_db, liftoff_db, m_id_trans_dict,
                                   miniprot_db, tree_dict, tgt_fai, ref_proteins, ref_trans, features_dict)
    else:  # processing features with exons
        if ENTRY_FEATURE:
            ref_trans_id = ref_gene_id
        else:
            ref_gene_id, ref_trans_id = lifton_utils.get_ref_ids_liftoff(features_dict,
                                                                         lifton_gene.entry.id, locus.id)
        lifton_trans, cds_num = lifton_add_trans_exon_cds(lifton_gene, locus, ref_db, liftoff_db, ref_trans_id)
        if cds_num > 0:
            liftoff_aln = LiftOn_liftoff_alignment(lifton_trans, locus, ref_proteins)
            miniprot_aln, valid = LiftOn_miniprot_alignment(locus, m_id_trans_dict, miniprot_db, ref_proteins)
            if liftoff_aln.identity < 1 and valid:
                cds_list = chaining_algorithm(liftoff_aln, miniprot_aln, tgt_fai)
                lifton_gene.update_cds_list(lifton_trans.entry.id, cds_list)
        lifton_gene.orf_search_protein(lifton_trans.entry.id, ref_trans_id, tgt_fai, ref_proteins, ref_trans)
    return lifton_gene


# Iterate through Liftoff features, pair them with corresponding miniprot transcripts, and run the protein-
maximization algorithm.
for feature in features:
    for locus in liftoff_db.features_of_type(feature):
        lifton_gene = protein_maximization(None, locus, ref_db.db_connection, liftoff_db, m_id_trans_dict,
                                  miniprot_db, tree_dict, tgt_fai, ref_proteins, ref_trans, features_dict, True)
```

### **Algorithm S2:** Mapping CDS boundaries onto a protein alignment

```
# Inputs:
#   - cds_lens (list of integers): Lengths of coding sequences.
#   - cds_protein_aln_boundary (list): List of tuples, each representing the start and end boundary of a CDS in
#     protein coordinates. - (cds_start, cds_end)
#   - cigar_ls (list): List of tuples, each representing the length and type - (cigar_len, cigar_symbol)

# Map the CDS boundaries onto proteins
function get_cds_protein_boundary(cds_lens):
    cds_cumulative = [sum(cds_lens[:i+1]) for i in range(len(cds_lens))]
    cds_cumulative_div = [x / 3 for x in cds_cumulative]
    cds_protein_boundary = {}
    for idx in range(len(cds_cumulative_div)):
        start = cds_cumulative_div[idx - 1] if idx > 0 else 0
        end = cds_cumulative_div[idx]
        cds_protein_boundary[idx] = (start, end)
    return cds_protein_boundary


# Adjust the CDS boundaries on protein-to-protein alignments.
function adjust_cds_protein_boundary(cds_protein_aln_boundary, D_accum_len, length):
    cds_boundary_shift = 0
    for i, (cds_start, cds_end) in enumerate(cds_protein_aln_boundaries):
        if (cds_start <= D_accum_len) and (cds_end >= D_accum_len):
            # Adjust CDS boundaries
            cds_boundary_shift += length
            cds_end += length
        cds_protein_aln_boundary[i] = (cds_start, cds_end)
    return cds_protein_aln_boundary


# Map CDS boundaries onto a protein alignment
cds_protein_boundary = get_cds_protein_boundary(cds_lens)
D_accum_len = 0
cds_protein_aln_boundary = cds_protein_boundary.copy()
for length, symbol in cigar_ls:
    if symbol == "D":
        # Deletion in CIGAR string => (longer protein sequence in the extract Liftoff or miniprot proteins)
        cds_protein_aln_boundary = adjust_cds_protein_boundary(cds_protein_aln_boundary, D_accum_len, length)
    D_accum_len += length
```

**Algorithm S3:** The chaining algorithm

```
# Inputs:
#   - liftoff_aln: An instance of Lifton_Alignment class, storing Liftoff parasail protein alignment information
#   - miniprot_aln: An instance of Lifton_Alignment, storing parasail miniprot parasail protein alignment information
#   - tgt_fai: An Fasta instance from the pyfaidx package, storing target genome sequences

# The main LiftOn chaining algorithm.
function chaining_algorithm(liftoff_aln, miniprot_aln, tgt_fai):
    l_children, m_children = liftoff_aln.cds_children, miniprot_aln.cds_children
    m_c_idx, l_c_idx, m_c_idx_last, l_c_idx_last = 0, 0, 0, 0
    cds_list, ref_aa_liftoff_count, ref_aa_miniprot_count, chains = [], 0, 0, []

    while m_c_idx != (length(m_children) - 1) or l_c_idx != (length(l_children) - 1):
        if m_c_idx == length(m_children) - 1 and l_c_idx < (length(l_children) - 1):
            l_c_idx, ref_aa_liftoff_count = push_cds_idx(l_c_idx, liftoff_aln, ref_aa_liftoff_count)
        else if m_c_idx < length(m_children) - 1 and l_c_idx == (length(l_children) - 1):
            m_c_idx, ref_aa_miniprot_count = push_cds_idx(m_c_idx, miniprot_aln, ref_aa_miniprot_count)
        else:
            m_c, l_c = m_children[m_c_idx], l_children[l_c_idx]
            if ref_aa_liftoff_count < ref_aa_miniprot_count:
                l_c_idx, ref_aa_liftoff_count = push_cds_idx(l_c_idx, liftoff_aln, ref_aa_liftoff_count)
            else if ref_aa_liftoff_count > ref_aa_miniprot_count:
                m_c_idx, ref_aa_miniprot_count = push_cds_idx(m_c_idx, miniprot_aln, ref_aa_miniprot_count)
            else:
                if l_c_idx > 0 and m_c_idx > 0 and m_c.end == l_c.end:
                    cdss = process_m_l_children(m_c_idx, m_c_idx_last, miniprot_aln, l_c_idx, l_c_idx_last,
                                                liftoff_aln, tgt_fai, chains)
                    cds_list += cdss
                    m_c_idx_last, l_c_idx_last = m_c_idx, l_c_idx
                l_c_idx, ref_aa_liftoff_count = push_cds_idx(l_c_idx, liftoff_aln, ref_aa_liftoff_count)
                m_c_idx, ref_aa_miniprot_count = push_cds_idx(m_c_idx, miniprot_aln, ref_aa_miniprot_count)
    l_c_idx, m_c_idx = l_c_idx + 1, m_c_idx + 1
    cds_list += process_m_l_children(m_c_idx, m_c_idx_last, miniprot_aln, l_c_idx, l_c_idx_last, liftoff_aln, tgt_fai,
chains)
    return cds_list, chains


# Calculate the accumulated amino acids in the alignments and group the CDSs together for processing.
function push_cds_idx (c_idx, lifton_aln):
    aa_start = 0
    aa_end = lifton_aln.cdss_protein_aln_boundaries[c_idx][1]
    aa_end = ceil(aa_end)
    ref_count = 0  # Calculate the accumulated amino acids in the reference protein alignment
    for i, letter in enumerate(lifton_aln.ref_seq[aa_start:aa_end]):
        if letter != "-":
            ref_count += 1
    c_idx += 1
    return ref_count, c_idx


# Process the grouped Liftoff and miniprot CDSs
function process_m_l_children(m_c_idx, m_c_idx_last, miniprot_aln, l_c_idx, l_c_idx_last, liftoff_aln, tgt_fai,
chains):
    m_aa_start, m_aa_end = get_protein_boundary(miniprot_aln.cdss_protein_aln_boundaries, m_c_idx_last, m_c_idx)
    l_aa_start, l_aa_end = get_protein_boundary(liftoff_aln.cdss_protein_aln_boundaries, l_c_idx_last, l_c_idx)
    m_matches, m_length = get_partial_id_fraction(miniprot_aln.ref_aln, miniprot_aln.query_aln, floor(m_aa_start),
ceil(m_aa_end))
    l_matches, l_length = get_partial_id_fraction(liftoff_aln.ref_aln, liftoff_aln.query_aln, floor(l_aa_start),
ceil(l_aa_end))
    cds_ls = create_lifton_entries(m_c_idx, m_c_idx_last, miniprot_aln, l_c_idx, l_c_idx_last, liftoff_aln, tgt_fai,
m_matches / m_length > l_matches / l_length)
    chains.append(if m_matches / m_length > l_matches / l_length then "miniprot" else "Liftoff")
    return cds_ls
```

**Algorithm S4:** Calculating protein and DNA sequence identity score

```
# Inputs:
#   - reference (string): The traceback reference alignment result.
#   - target (string): The traceback target alignment result.
#   - start (int): 0-based index of the start of the protein segment.
#   - end (int): 0-based index of the end of the protein segment.

# Calculate the partial protein sequence identity score for chaining algorithm
function get_partial_id_fraction(reference, target, start, end):
    matches, gaps_in_ref = 0, 0
    for i, letter in enumerate(reference[start:end]):
        if letter == '-':
            gaps_in_ref += 1
        if letter == target[i + start]:
            matches += 1
        if target[i + start] == "*":
            break
    total_length = (end - start) - gaps_in_ref # Longer protein without premature stop codon is not penalized
    if total_length == 0:
        return matches, 1
    return matches, total_length


# Calculate the full-length protein sequence identity score
function get_AA_id_fraction(reference, target):
    matches, gaps_in_ref = 0, 0
    for i, letter in enumerate(reference):
        if letter == '-':
            gaps_in_ref += 1
        if letter == target[i + start]:
            matches += 1
        if target[i + start] == "*":
            break
    if max(len(reference), len(target)) == 0:
        return matches, 1
    # Gap-compressed sequence identity
    total_length = max(len(reference), len(target)) - gaps_in_ref
    return matches, total_length


# Calculate the full-length transcript DNA sequence identity score
function get_DNA_id_fraction(reference, target):
    matches = 0
    # BLAST identity
    for i, letter in enumerate(reference):
        if letter == target[i]:
            matches += 1
    if max(len(reference), len(target)) == 0:
        return matches, 1
    return matches, max(len(reference), len(target))
```
